# Rifampicin-induced *Staphylococcus aureus* persister formation is driven by CodY regulon and oxidative stress level

**DOI:** 10.64898/2026.03.03.709237

**Authors:** Nicola Pordone, Johann Guillemot, Killian Rodriguez, Nicolas Personnic, Elisabeth Botelho-Nevers, Paul O. Verhoeven

## Abstract

*Staphylococcus aureus* is a major human pathogen. Beyond resistance, antibiotic persistence and tolerance reduce treatment efficacy. Persisters are characterized by heterogenous sub-populations of non-growers within an otherwise susceptible replicative population. We leveraged the spectral properties of Timer and Timer^FAST^—two novel growth reporter fluorescent proteins in *S. aureus*—to identify non-growers and population heterogeneity using live-cell microscopy and flow cytometry. Single-cell phenotype analysis during rifampicin exposure revealed population heterogeneity, with replicating cells coexisting alongside non-growing but viable cells. The non-grower phenotype increased in the *codY* mutant upon rifampicin treatment. We uncovered that *codY* mutant exhibits higher ROS level than the wild-type. Treatment with both rifampicin and menadione—a ROS inducer—triggered a fully non-growing population independently of CodY. These findings highlight the major role of CodY regulon and oxidative stress in persister formation and pave the way to develop new therapeutic strategies.

## INTRODUCTION

*Staphylococcus aureus* remains a leading cause of death from bacterial infections worldwide [1]. It colonizes approximately two billion individuals, causing both mild and life-threatening infections [2]. Healthy carriers face higher risks of self-infection, and while decolonization reduces this risk, failures or rapid recolonization are common despite *S. aureus* being susceptible to antimicrobials [3,4]. Belonging to the ESKAPE group of multidrug-resistant pathogens, *S. aureus* urgently requires novel antimicrobial strategies [5]. Interestingly, while antibiotic resistance complicates treatment, the rise in *S. aureus* infections over the past decade has coincided with a decrease in methicillin resistance [6].

Beyond resistance, *S. aureus* can survive antibiotics through adaptative mechanisms [7–9]. Antibiotic-tolerant cells exhibit an increased minimum duration for killing and therefore survive treatment longer [10]. Persisters are defined as a subpopulation of non-replicating bacteria, within a susceptible population, that tolerates antibiotics or other stresses through mechanisms that are not genetically driven [10,11]. Both tolerant and persistent cells resume growth once treatment ends [10,11]. Persistence constitutes a population-level phenomenon that implies phenotypic heterogeneity within the bacterial population [10]. Together, persistence and tolerance, collectively termed ’recalcitrance’, contribute to treatment failure but remain less studied than resistance [12].

Antibiotic exposure and environmental stresses such as phagocytes environment and nutrient deprivation have been shown to induce persistence and tolerance in *S. aureus* [9,13–15]. Acidic environment promoted non-stable small colonies, considered hallmarks of persisters [9,16–18]. Macrophage-derived reactive nitric and oxygen species (RNS/ROS) collapsed *S. aureus* metabolic activity, leading to enhanced persistence [14,15]. The stringent response, a (p)ppGpp-mediated alarmone system that enables adaptation to stress and nutrient deprivation, has been shown to drive *S. aureus* persister formation within antibiotic-treated macrophages [13]. Remarkably, in planktonic culture, persister formation upon antibiotic exposure occurred independently of this pathway [8]. Thus, the molecular mechanisms underlying *S. aureus* persistence remains incompletely understood.

Recently, Helaine and colleagues highlighted the importance of single-cell approaches in persister studies [12]. Although biphasic killing curves from time-kill assays are widely used to identify persisters, the method is labor-intensive and lacks single-cell resolution, driving the development of microfluidic and fluorescent-based approaches [19]. The Timer, a fluorescent protein (FP) that shifts from green to red over time, has proven to be a powerful tool for tracking, isolating, and analyzing persisters at the single-cell level in *Legionella* and *Salmonella* [20–22]. In *S. aureus*, a couple of fluorescence-based approaches have been developed [8,13,23], but Timer as not yet been implemented. To fill this gap, we engineered Timer and Timer^FAST^, a fast-maturating variant, optimized for *S. aureus,* that enable monitoring of *S. aureus* replication dynamics in real-time using green-to-red fluorescence ratio as a proxy of growth rates.

Using the spectral properties of Timer^FAST^ in *S. aureus*, we investigated how the stringent response drives persister formation following antibiotic stress. In *S. aureus*, the CodY regulon contributes to the stringent response by derepressing target genes under amino acid starvation and low GTP conditions [13]. To elucidate the interplay between the stringent response and persistence, we used rifampicin, a clinically relevant bactericidal antibiotic commonly employed against *S. aureus*. While rifampicin resistance mechanisms are well defined, the dynamics of persister formation in response to this drug remain largely unknown.

Here, we demonstrated that rifampicin exposure induces persisters alongside resistant cells. We further found that rifampicin triggers ROS accumulation, a condition strongly linked to the persister phenotype. Strikingly, ROS-associated persister formation was amplified in *codY* mutant, highlighting the role of the CodY regulon in modulating oxidative stress and persistence. Together, our findings provide new insights into the molecular mechanisms of *S. aureus* persistence upon rifampicin treatment and establish a novel connection between CodY regulon, oxidative stress, and persister cell formation.

## RESULTS

### Timer and Timer^FAST^ engineering in *S. aureus*

In our previous work, we used DsRed, which exhibits slow chromophore maturation of approximately 27 hours [24–26]. This property caused DsRed to accumulate in non-growing bacteria, enabling tracking of replication dynamics inside eukaryotic cells via fluorescence dilution [26]. However, fast-replicating bacteria lost fluorescence within a few hours, limiting detection of rapidly dividing populations [26].

To overcome these limitations, we performed site-directed mutagenesis on the *DsRed* gene to introduce the S197T substitution, thereby generating a Timer FP identical to that originally described [27]. As expected, Timer-expressing *S. aureus* grown on agar plate exhibited fluorescence shifting from green to red over time (Fig. 1a, Extended Data Fig. 1b,c,d). At 48 hours, Timer green fluorescence was comparable to GFP, but at 24 hours it was markedly weaker, making colony detection more challenging (Fig. 1a, Extended Data Fig. 1a,b). We reasoned that the weak signal at 24 hours may result from fluorescence dilution of the green-emitting chromophore (Fig 1a, Extended Data Fig. 1a,b), as previously reported for DsRed [26]. We then used live-cell confocal microscopy to further assess Timer behavior in fast-replicating *S. aureus* (Fig. 1b,c). As expected, green fluorescence progressively diluted during microcolony formation, confirming that Timer maturation lagged behind bacterial doubling time (Fig 1c). To accelerate chromophore maturation, we tested mutations previously described in DsRed [28]. The most promising candidate, exhibiting S197T, N42Q and V105A mutations, was called Timer^FAST^. Strikingly, Timer^FAST^-expressing *S. aureus* grown in rich medium retained strong green fluorescence up to 7 hours, whereas Timer-expressing bacteria rapidly lost green signal (Fig 1c, Extended Data Fig. 1e, Supplementary Videos 1 and 2). In addition to faster maturation, Timer^FAST^was also 2 to 3 times brighter than those of Timer (Extended Data Fig. 1b,e). For both proteins, growth arrest was correlated with red fluorescence emission (Fig. 1a, d, Extended Data Fig. 1c,f, Supplementary Videos 3 and 4). While the red fluorescence intensity was comparable, the red chromophore also maturated faster in Timer^FAST^ (Fig. 1a,d, Extended Data Fig. 1f). Timer^FAST^ displayed a rapid green-to-red fluorescence shift, allowing detection of growth arrest as early as 1 hour after starvation onset (Fig. 1e). In contrast, Timer shifted more slowly, between 4 and 8 hours (Fig. 1e), making it suitable for studying slow-growing bacteria. To further characterize Timer and Timer^FAST^, we performed flow cytometry. Bacteria were grown in rich medium (TSB90), poor medium (TSB5), or in HBSS to induce fast-, slow- and non-growing populations, respectively (Extended Data Fig. 1g). Mean doubling times were 24 and 23 min for fast-growing, 91 and 87 min for slow-growing, and >1440 min for non-growing bacteria expressing Timer and Timer^Fast^, respectively (Fig. 1f). In HBSS, the mean log(G/R) ratios were near zero, with slightly higher values for Timer^FAST^ due to its stronger high green fluorescence (Fig. 1g, Extended Data Fig. 1h,i). Notably, only Timer^FAST^ clearly distinguished fast- and slow-growing populations, with log(G/R) ratios of 1.32 ± 0.07 and 1.07 ± 0.10, respectively (Fig. 1g, Extended Data Fig. 1h,i). Flow cytometry confirmed that Timer^FAST^ accurately detects subtle changes in generation time under fast-growing conditions, while Timer, with slower maturation, is better suited for extended generation times. Collectively, the complementary dynamics of Timer and Timer^FAST^ broaden the monitoring of bacterial growth states.

**Fig. 1.**
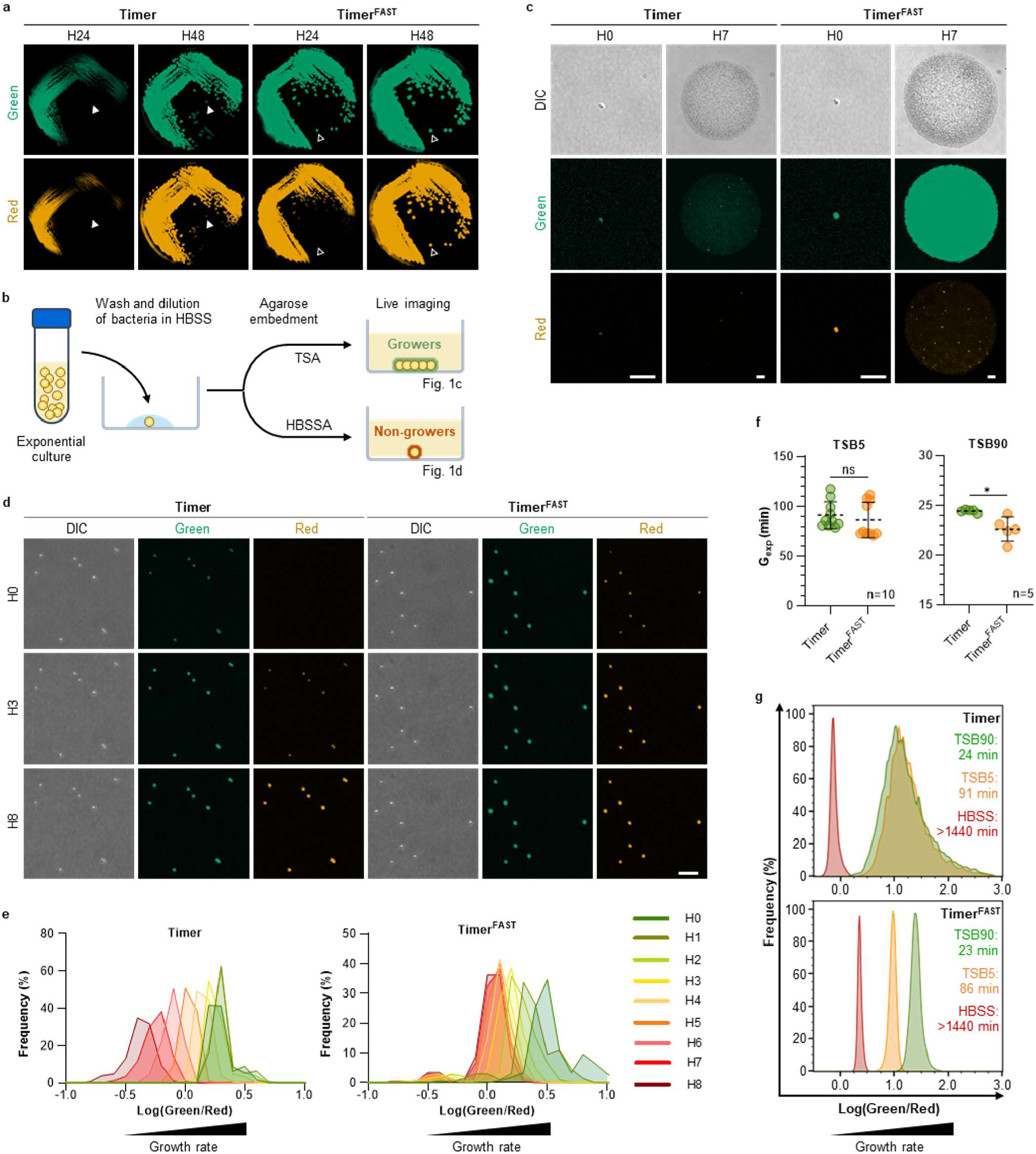
Spectral dynamics of *S. aureus* expressing Timer and Timer^FAST^. **a**, Colonies of *S. aureus* SF8300 constitutively expressing Timer or Timer^FAST^ grown for 24 and 48 hours on TSA plates supplemented with chloramphenicol. Open and closed arrowheads indicate the same colony tracked over time. **b**, Experimental workflow for live-cell imaging of *S. aureus* microcolonies. **c–d**, Live-cell confocal microscopy of Timer- and Timer^FAST^-expressing *S. aureus* grown for 7 hours in TSA (c) or for 8 hours in HBSSA (d). Images correspond to maximum intensity projections in z. Green channel: Ex 488 nm, Em 525/50 nm. Red channel: Ex 561 nm, Em 600/52 nm. Scale bar: 10 µm. **e**, Distribution of single-cell fluorescence ratios from three independent biological experiments illustrated in (d) (n = 223 for Timer at H0, n = 260 for Timer^FAST^ at H0). **f**, Generation times during exponential growth (G_exp_) of Timer- and Timer^FAST^-expressing *S. aureus* grown in TSB90 (9:1 TSB:HBSS) or TSB5 (5:95 TSB:HBSS). Each dot represents a value calculated from OD measurements acquired from a single well across three independent biological experiments. Dotted lines and errors bars: mean ± SD. Mann-Whitney U test: * *P* < 0.05; ns, not significant. **g**, Fluorescence ratios of Timer- and Timer^FAST^-expressing *S. aureus* grown in TSB90, TSB5, or HBSS. Data are from one representative experiment out of three independent biological replicates. G_exp_ values are indicated for each condition. DIC: differential interference contrast. HBSS: Hank’s balanced salt solution. HBSSA: Hank’s balanced salt agar. TSA: tryptic soy agar. TSB: tryptic soy broth.

### Rifampicin persisters emerge alongside resistant *S. aureus* following rifampicin treatment *in vitro*

Rifampicin is a broad-spectrum antibiotic commonly used in the treatment of *S. aureus* infections [29]. It exerts its bactericidal activity by binding to the β-subunit of RNA polymerase (RpoB), thereby inhibiting transcription [30,31]. In *S. aureus*, resistant mutants can arise spontaneously, as rifampicin exposure during the exponential growth induces mutation in the *rpoB* gene [8].

To investigate the dynamics of *S. aureus* survival under antibiotic pressure, we first performed time-kill experiments (Fig. 2a). Using rifampicin at 10× the minimum inhibitory concentration (MIC), we observed a clear bactericidal effect by 18 hours, followed by pronounced regrowth at 24 hours, a hallmark of emerging resistance (Fig. 2b).

**Fig. 2.**
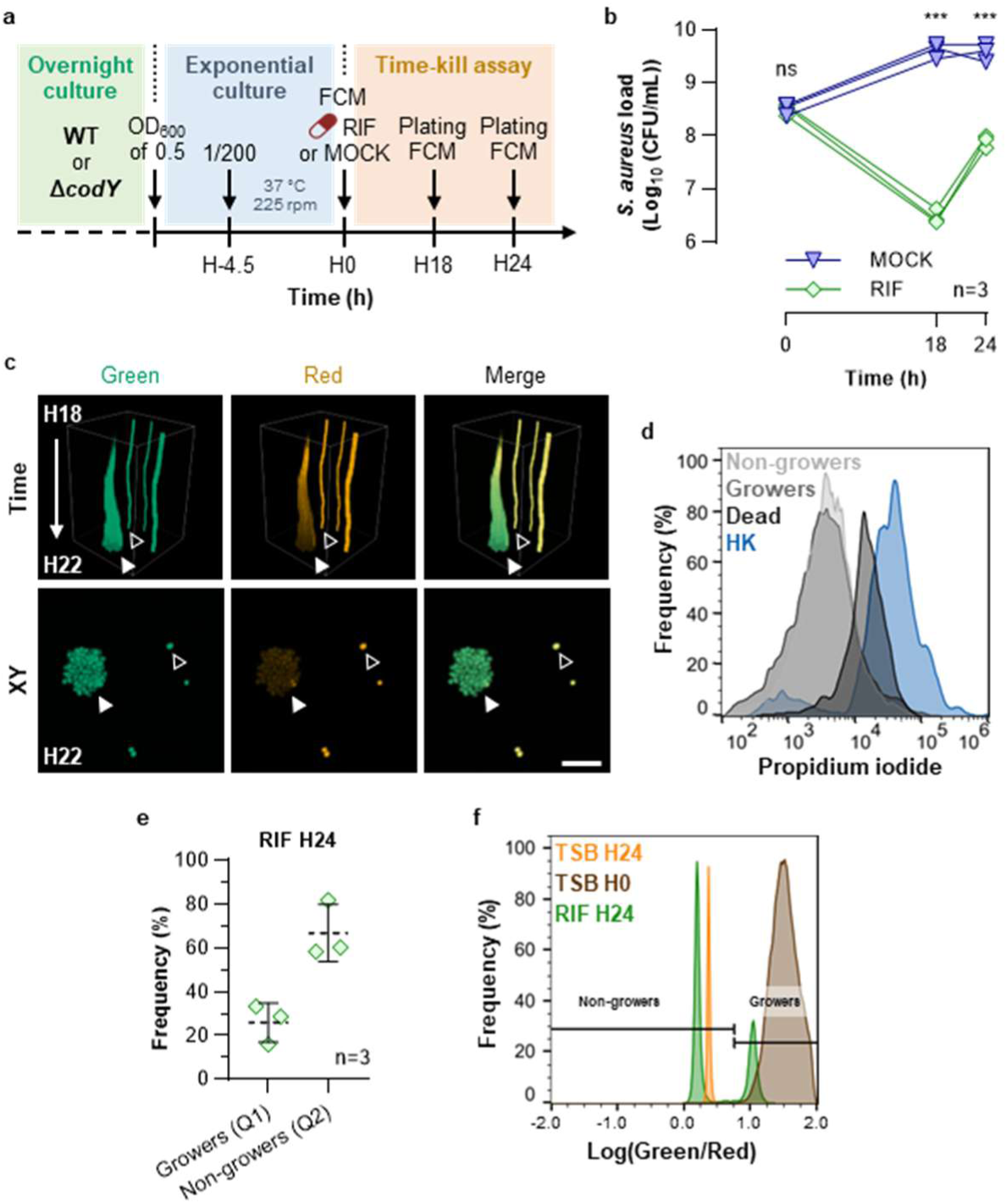
Emergence of *S. aureus* persisters and resistant cells during rifampicin treatment. **a**, Experimental workflow of Timer^FAST^-expressing *S. aureus* HG001 WT treated with 10x minimum inhibitory concentration (MIC) rifampicin and analyzed by culture (Fig 2b), live-cell confocal microscopy (LCM, Fig 2c), and flow cytometry (FCM, Fig 2d-f). **b**, Time-kill assay of Timer^FAST^-expressing *S. aureus* HG001 WT treated with 10x MIC rifampicin (RIF) or DMSO (MOCK). A curve represents an independent biological replicate (n=3). Two-way ANOVA: \*\*\**P* value < 0.001 (RIF vs. MOCK). **c**, Live-cell confocal microscopy of Timer^FAST^-expressing *S. aureus.* Bacteria embedded in TSA supplemented with 10x MIC rifampicin were imaged from 18 (H18) to 22 (H22) hours. Time shows 3D maximum intensity projection over time. XY shows Z-maximum intensity projection at H22. Open arrowheads: non-growers. Close arrowheads: growers. Green channel: Ex 488 nm, Em 525/50 nm. Red channel: Ex 561 nm, Em 600/52 nm. Scale bar: 10 µm. d, Propidium iodide (PI) uptake in Timer^FAST^-expressing *S. aureus* after 24 hours of 10xMIC rifampicin treatment. Growers (Q1), non-growers (Q2) and dead cells (Q3) correspond to the gating strategy shown in Extended Data Fig.2a. Heat-killed cells, used as PI-positive controls, were not exposed to rifampicin. Data shown are from one representative of three independent flow cytometry experiments. **e**, Frequency of growers (Q1) and non-growers (Q2) in Timer^FAST^-expressing *S. aureus* after 24 hours of 10xMIC rifampicin treatment, measured by flow cytometry. Gating strategy is shown in Extended Data Fig.2a. Each dot represents an independent experiment. Dotted lines and error bars: mean ± SD. **f**, Fluorescence ratios in Timer^FAST^-expressing *S. aureus* grown in exponential (TSB H0), in stationary phase (TSB H24), or treated 24 hours with 10x MIC rifampicin (RIF 24H). Data shown are from one representative of three independent flow cytometry experiments.

Live single-cell confocal microscopy at 18 hours using Timer^FAST^-expressing *S. aureus* highlighted phenotypic heterogeneity with non-growers co-existing alongside rifampicin-resistant mutants under antibiotic pressure (Fig. 2c). To further investigate these populations, we performed flow cytometry on Timer^FAST^-expressing *S. aureus* after 24 hours of rifampicin treatment (Fig. 2d-f, Extended Data Fig. 2a-c). Gating strategy was established using growers (Q1), non-growers (Q2) and heat-killed bacteria (Q3) (Extended Data Fig. 2a). Consistent with microscopy, Timer^FAST^ spectral analysis showed extensive growth rate heterogeneity at the single-cell level. Interestingly, the most abundant population was non-growers (Fig. 2e), which retained green fluorescence and excluded propidium iodide (Fig. 2d), indicating maintained metabolic activity and intact cell envelopes during rifampicin exposure. In contrast, a subpopulation underwent cell death, characterized by loss of Timer^FAST^ fluorescence and propidium iodide uptake (Q3) (Fig. 2d). We also observed a subpopulation of replicating bacteria corresponding to rifampicin-resistant mutants (Q1) that emerged under antibiotic pressure (Fig. 2f, Extended Data Fig.2a).

In line with recent clarifications on antibiotic persistence [10], these results collectively demonstrate that rifampicin exposure generates a heterogeneous *S. aureus* population, including a subpopulation of rifampicin-tolerant cells that fulfill the criteria for persisters.

### CodY regulon promotes *S. aureus* survival and growth rate during rifampicin treatment

To investigate the molecular mechanisms underlying persister formation during rifampicin treatment, we performed time-kill assays (Fig 3a) using several *S. aureus* mutant strains deficient in genes involved in the stringent response (data not shown). We focused on the *codY* mutant, which displayed a tenfold lower bacterial burden at 18 hours and struggled to regrow at 24 hours compared to the WT strain (Fig. 3a,b). This regrowth defect was no longer observed by 48 hours, indicating that rifampicin-resistant mutants had fully repopulated both cultures (Fig. 3a,b). To explore the basis of the regrowth defect, we assessed the fitness of the *codY* mutant. Optical density measurements at 600 nm (OD_600nm_) showed a longer generation time for the *codY* mutant (Extended Data Fig. 3a). However, direct quantification of CFUs during exponential growth revealed higher CFUs for the *codY* mutant (Extended Data Fig. 3b), ruling out an intrinsic fitness defect. As previously reported [32], the *codY* mutant readily forms aggregates (Extended Data Fig. 3c), which decrease OD-based measurements and likely explain the discrepancy between OD values and CFU counts.

**Fig. 3.**
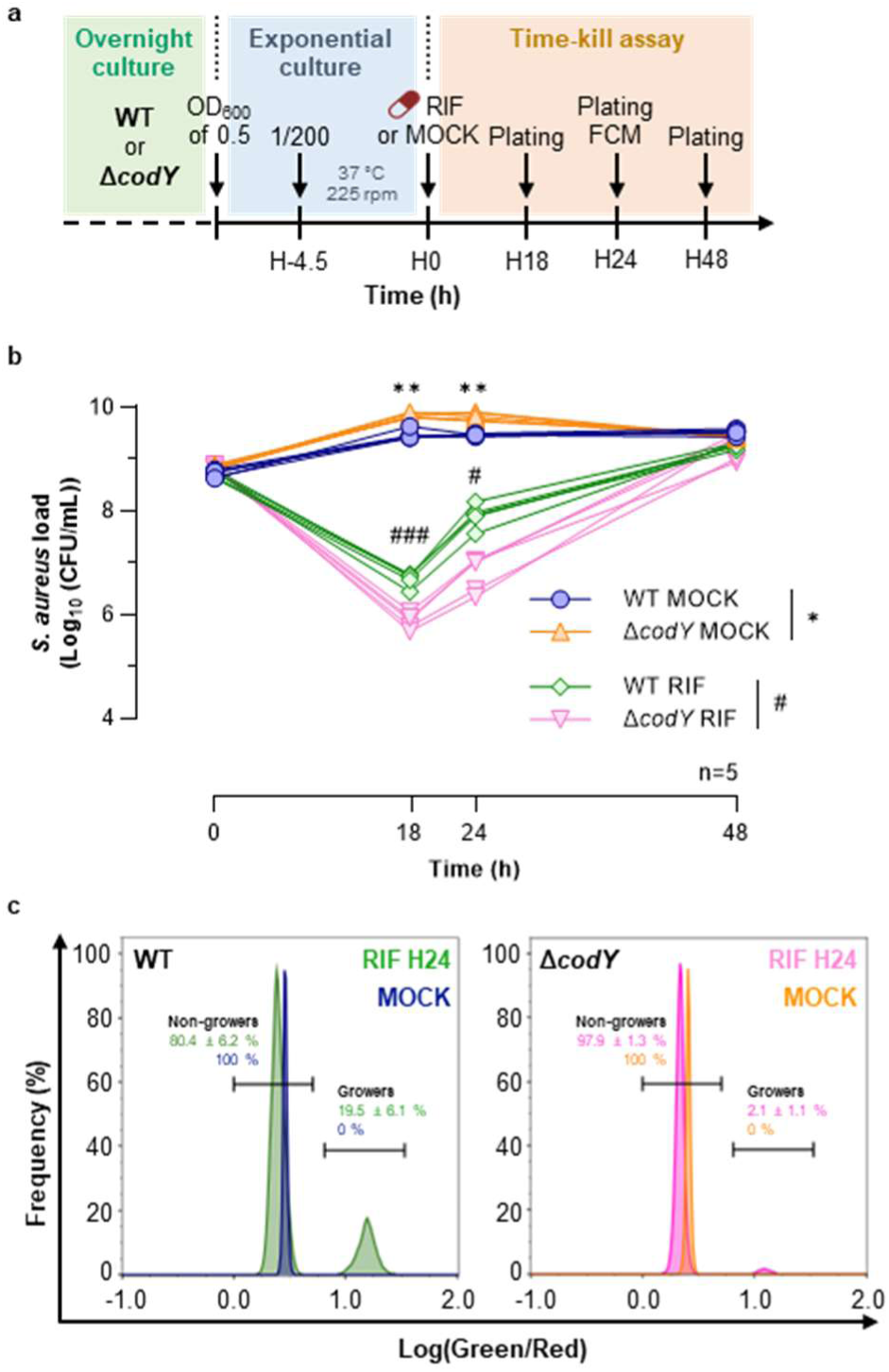
CodY regulon promote *S. aureus* survival and growth rate during rifampicin treatment. **a**, Experimental workflow of Timer^FAST^-expressing *S. aureus* HG001 WT and *codY* mutant strains treated with 10x minimum inhibitory concentration (MIC) rifampicin (RIF) and analyzed by culture (Plating, b) and flow-cytometry (FCM, c). **b**, Time-kill assay of Timer^FAST^*-*expressing *S. aureus* WT and *codY* mutant strains treated with 10x MIC rifampicin (RIF) or DMSO (MOCK) for 48 hours. Each curve represents an independent biological replicate. Two-way ANOVA: * or ^#^ P value < 0.05; ** or ^##^ P value < 0.01; *** or ^###^ P value < 0.001. **c-d**, Fluorescence ratios of Timer^FAST^-expressing *S. aureus* WT (left) and *codY* mutant strain (right) after 24 hours treatment with DMSO (MOCK) or 10x MIC RIF. Data shown are from one representative of five independent flow cytometry experiments. Frequencies (mean ± SD calculated from the five experiments) are indicated for each condition.

In flow cytometry, non-growers represented the most abundant population in both strains. However, the proportion of growing cells was markedly reduced in the *codY* mutant (Fig. 3c). We next assessed the rifampicin mutation rate of WT and *codY* mutant strains (Extended Data Fig. 3d). Across three replicates, only one resistant colony emerged from the *codY* mutant, corresponding to a mean mutation ratio of 1.9 × 10^-9^, whereas multiple colonies arose from WT, with a mean ratio of 1.9 × 10^-8^ (Extended Data Fig. 3d). These results indicate that CodY acts as a pro-growth factor, promoting rifampicin resistance associated mutations and/or supporting bacterial growth during rifampicin exposure.

To further examine the pro-growth role of CodY during rifampicin treatment, bacteria in the exponential growth phase were treated with rifampicin for 18 hours and embedded in TSA (with or without rifampicin) for live-cell confocal microscopy. Images were acquired every 20 min between 18 and 21 hours (Fig. 4a, Extended Data Fig. 4a). We used 100× MIC to maximize the likelihood of imaging microcolonies originating from single cells. At 18 hours, *S. aureus* microcolony volumes were higher in the *codY* mutant than in WT (Fig. 4b,d), likely due to aggregates formed by the mutant (Extended Data Fig. 3c), which were difficult to fully dissociate before embedding in TSA. Strikingly, we observed that *codY* mutant struggled to resume growth after 18 hours of rifampicin treatment compared to WT strain (Fig. 4b,c,d,g). When pre-exposed to rifampicin, only the WT strain formed large microcolonies at 21 hours, even when embedded in rifampicin-free TSA after 18 hours (Fig. 4b,c,g). Mock-treated WT and *codY* mutant strains formed large microcolonies between 18 and 21 hours in TSA but not in TSA with rifampicin (Extended Data Fig. 4b,c,d,g), reflecting that stationary-phase cells readily form rifampicin-tolerant populations [8]. Thus, we analyzed the G/R fluorescence ratio of Timer^FAST^-expressing *S. aureus* during microcolony formation between 18 and 21 hours after rifampicin treatment, with or without rifampicin selection in the embedding agar (Fig. 4d-i, Extended Data Fig. 4d-i). In both WT and *codY* mutant strains, mock-treated bacteria formed microcolonies composed of replicating cells, as indicated by Log(G/R) values progressively shifting toward 1 in the absence of rifampicin (Extended Data Fig. 4g-i). In rifampicin-treated WT cells, Log(G/R) values increased even with rifampicin selection, reflecting the emergence of resistant mutants (Fig. 4d,e,g,h). Notably, a small subpopulation of WT cells maintained Log(G/R) values near zero, indicating the presence of growth-arrested bacterial cells regardless of rifampicin selection (Fig. 4e,h). Such growth-arrested cells were also observed in mock-treated conditions, indicating that “spontaneous” growth-arrested bacterial cells can emerge within the population independently of rifampicin pre-exposure (Extended Data Fig. 4g-i). Strikingly, in the *codY* mutant, growth-arrested phenotype was predominant irrespective of rifampicin selection in the embedding agar (Fig. 4f,i). The inability of *codY* mutant to resume growth in the absence of rifampicin selection suggests that CodY is required for efficient recovery following rifampicin-induced stress. To assess whether growth-arrested CodY-deficient bacteria could resume growth after rifampicin treatment, overnight cultures were normalized to 8 Log(CFU/ml) (Fig. 4j), exposed to rifampicin at 100× MIC for 18 hours (Fig. 4k), washed in PBS, and serially diluted onto agar plates with or without rifampicin before 24-hour incubation (Fig 4l). The WT strain regrew on both media, confirming the emergence of rifampicin-resistant mutants (Fig. 4l). In contrast, the *codY* mutant failed to grow in the presence of rifampicin, consistent with a lower probability of resistant mutant emergence (Fig. 4l). Notably, since the *codY* mutant only resumed growth on TSA without rifampicin, the growth-arrested cells observed by live-cell microscopy likely correspond to persisters (Fig. 4l).

**Fig. 4.**
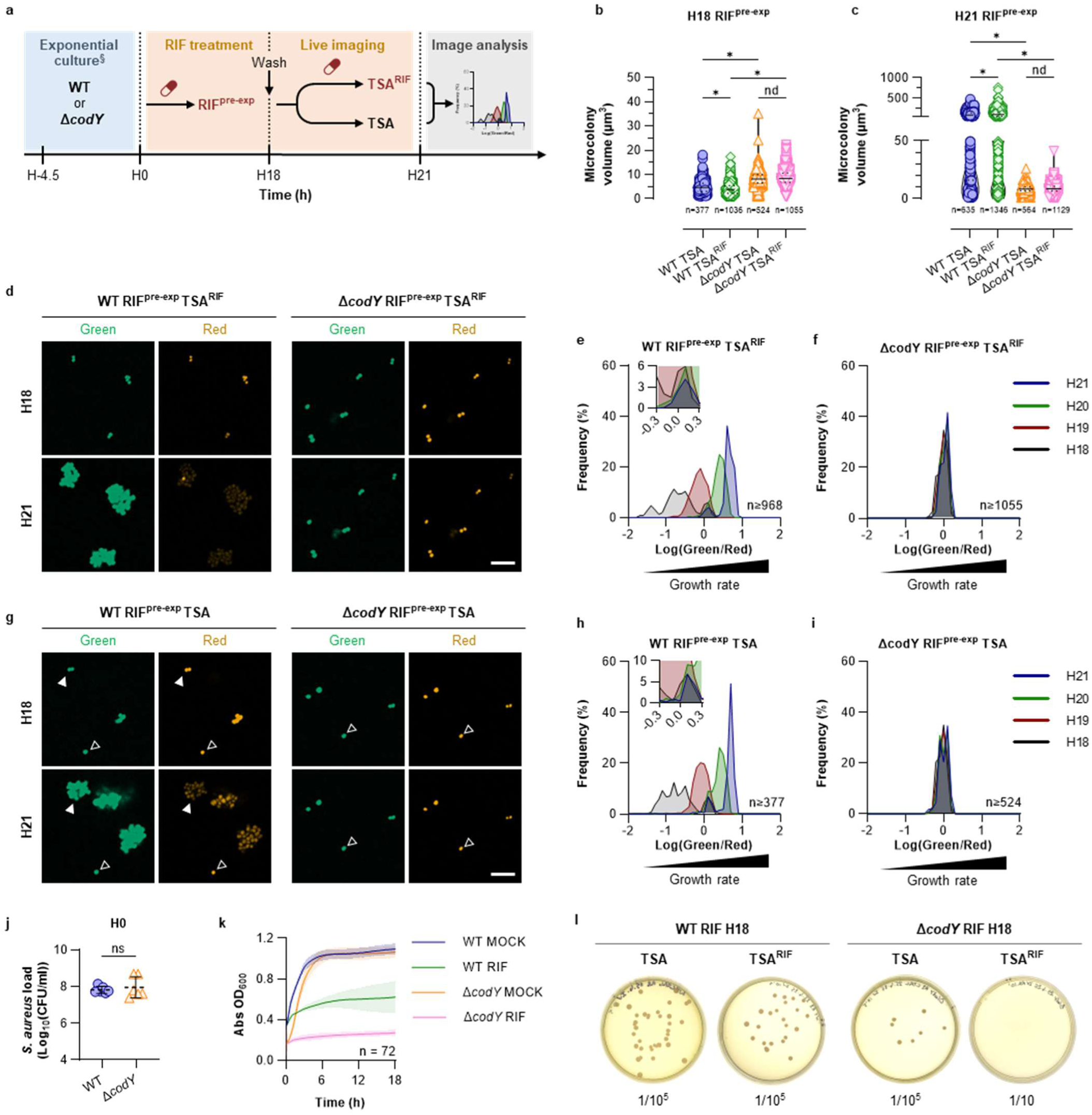
*codY* deficiency increase the non-growing population during rifampicin treatment. **a**, Experimental workflow of Timer^FAST^-expressing *S. aureus* HG001 wild-type (WT) and *codY* mutant strains treated with 100x minimum inhibitory concentration (MIC) rifampicin (RIF^pre-exp^) in TSB during 18 hours (H18), embedded in TSA or in TSA supplemented with 100x MIC rifampicin (TSA^RIF^), and analyzed by live-cell confocal microscopy (b-i). ^§^Exponential cultures were prepared as described in Fig. 3a. **b,c**, *S. aureus* microcolony volume at 18 (H18, b) and 21 (H21, c) hours. Values (n) indicate the number of microcolony measured across three independent biological experiments. Solid and dotted lines: median ± IQR. Kruskal-Wallis test with Benjamini and Hochberg false discovery rate correction for multiple comparisons: * Q < 0.05; nd, not discovered. **d,g**, Representative images of *S. aureus* microcolony embedded in TSA^RIF^ (d), or TSA (g), at 18 (H18) and 21 (H21) hours. Open arrowheads: non-growers. Close arrowheads: growers (rifampicin-resistant mutant). Green channel: Ex 488 nm, Em 525/50 nm. Red channel: Ex 561 nm, Em 600/52 nm. Scale bar: 10 µm. **e,f,h,i**, Green to red fluorescence ratios of *S. aureus* microcolonies from 18 (H18) to 21 (H21) hours. The number of microcolonies (n) analyzed across three independent biological experiments is indicated on the graph. Insets depict an enlarged view of the region of the x-axis near zero. **j**, Bacterial load of *S. aureus* HG001 wild-type or Δ*codY* strains grown for 4.5 hours in TSB prior to the experiment shown in (k) (corresponding to the H0 time point in Fig. 4a). Each dot represents the value obtained from a single broth across three independent biological experiments. Dotted lines and error bars: mean ± SD. Mann-Whitney U test: ns, not significant. **k**, Growth kinetics of *S. aureus* HG001 wild-type or Δ*codY* strains treated with either 100x MIC rifampicin or DMSO (MOCK) at t=0. The inocula quantified in (j) were used for each corresponding replicate. Each dot represents a single well acquired across three independent biological experiments. **i**, Culture on TSA and TSA^RIF^ of *S. aureus* HG001 wild-type or Δ*codY* strains following the experiment shown in (k), corresponding to 18 hours of treatment with 100x MIC rifampicin. Dilution factors of the inocula are indicated for each condition. Images are from a single experiment.

Together, our results indicate that CodY acts as a pro-growth factor that promotes regrowth after rifampicin stress, thereby enhancing the likelihood of resistance emergence.

### CodY regulates reactive oxygen species-mediated persistence during rifampicin exposure

Given that macrophage-derived RNS and ROS promote rifampicin tolerance [14] and that the *codY* mutant exhibited growth arrest under rifampicin, we investigated whether CodY regulates oxidative metabolism in *S. aureus* during rifampicin exposure.

During exponential phase, *S. aureus*-derived ROS level was higher in the *codY* mutant than in the WT, whereas stationary-phase measurements showed no difference (Fig. 5a,b), suggesting that CodY, in its GDP-bound active form, regulates oxidative metabolism. Cumulative measurement of ROS from lag (H0) to stationary phase (H18) confirmed that the *codY* mutant accumulated more ROS than the WT strain (Fig. 5c). Rifampicin increased ROS levels in both strains, but this effect was stronger in the *codY* mutant than the WT strain, with fold changes of 1.7 versus 1.2, respectively (Fig. 5c). These findings indicate that CodY prevents ROS accumulation during rifampicin stress. Since rifampicin is known to trigger ROS production in *S. aureus* [33], we hypothesized that the growth-arrested phenotype of the rifampicin-treated *codY* mutant was due to excessive intracellular ROS, caused by the combined effects of drug exposure and CodY deficiency [33,34].

**Fig. 5.**
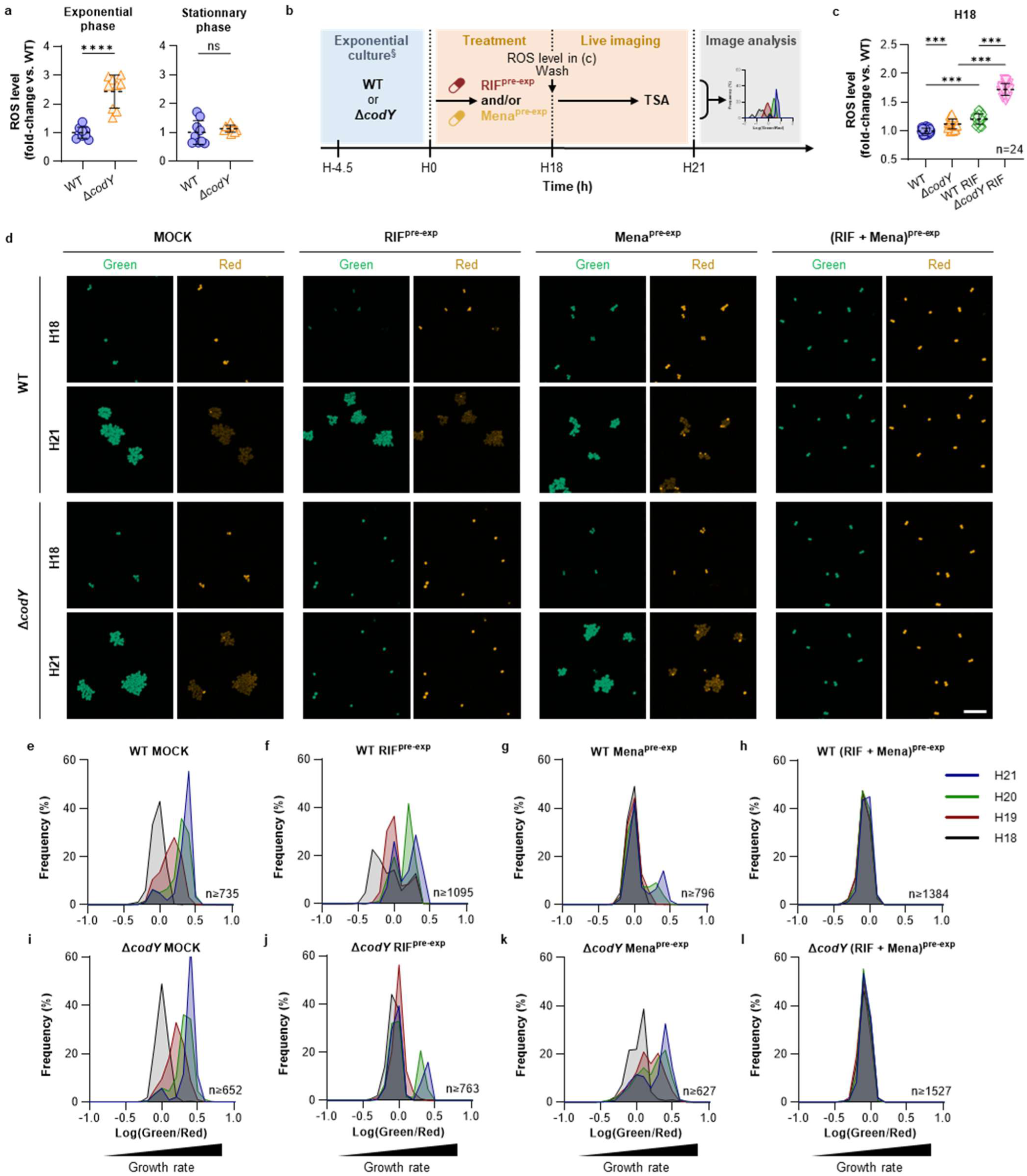
Reactive oxygen species promotes a non-growing phenotype of *S. aureus* during rifampicin treatment. **a**, Basal reactive oxygen species (ROS) accumulation in stationary and exponential phase in *S. aureus* HG001 wild-type (WT) and *codY* mutant strains. Each dot represents a single well acquired across three independent biological experiments. Dotted lines and error bars: mean ± SD. Mann-Whitney U test: *** P < 0.001; ns, not significant. **b**, Experimental workflow of Timer^FAST^-expressing *S. aureus* HG001 wild-type (WT) and *codY* mutant strains treated with DMSO (MOCK), 100x minimum inhibitory concentration (MIC) rifampicin (RIF^pre-exp^) or menadione (Mena) in TSB for 18 hours (H18), embedded in TSA, and analyzed by live-cell confocal microscopy (c-l). ^§^Exponential cultures were prepared as described in Fig. 3a. **c**, ROS level produced by *S. aureus* HG001 WT and *codY* mutant strains treated with DMSO (MOCK) or 100x MIC rifampicin during 18 hours. Each dot represents a single well acquired across six independent biological experiments. Dotted lines and error bars: mean ± SD. Mann-Whitney U test: *** P < 0.001. **d**, Representative images of *S. aureus* microcolony embedded in TSA at 18 (H18) and 21 (H21) hours. Open arrowheads: non-growers. Close arrowheads: growers (rifampicin-resistant mutant). Green channel: Ex 488 nm, Em 525/50 nm. Red channel: Ex 561 nm, Em 600/52 nm. Scale bar: 10 µm. **e-l**, Green to red fluorescence ratios of *S. aureus* microcolonies from 18 (H18) to 21 (H21) hours. The number of microcolonies (n) analyzed across three independent biological experiments is indicated on the graph. For each experiment, images were acquired every hour from 3 fields of view.

**Fig. 6.**
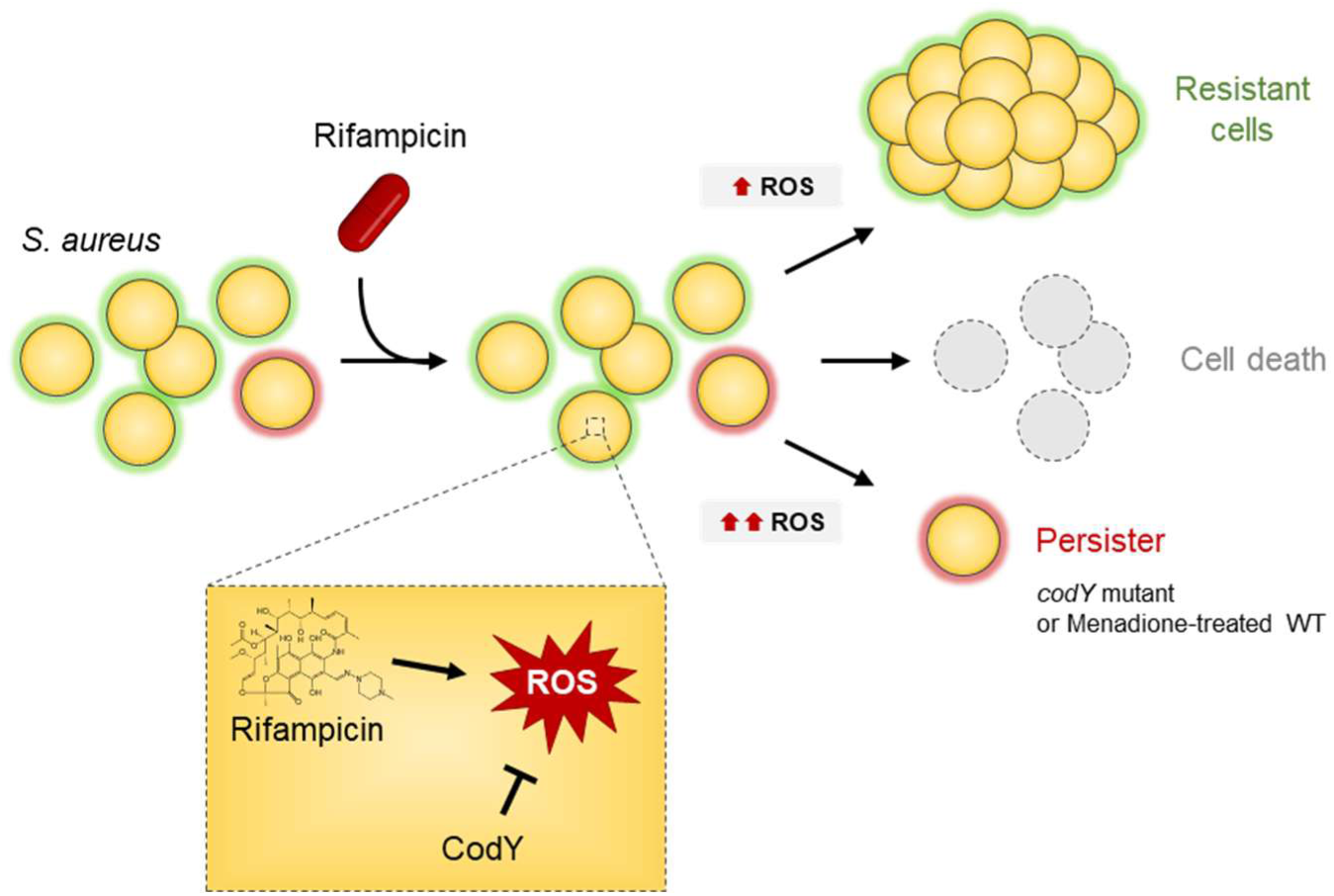
Model of CodY-dependent regulation of persister cell fate under rifampicin treatment. Upon exposure to rifampicin, *S. aureus* cells experience increased reactive oxygen species (ROS) production. Most cells either die or develop resistance, while a small sub-population enters a growth-arrested persister state, which likely correspond to the definition of persisters. Deletion of *codY* or treatment with menadione further enhances ROS accumulation, promoting the formation of a distinct population of persisters characterized by prolonged growth arrest and high rifampicin tolerance. CodY acts as a negative regulator of ROS levels under rifampicin stress. Schematic inset: rifampicin induces ROS production, which is modulated by CodY.

To test the impact of ROS independently of rifampicin, we exposed cells to menadione, a redox-cycling compound that elevates endogenous ROS in *S. aureus* [14] (Fig. 5b, Extended Data Fig. 5a). In the WT strain, Log(G/R) patterns with or without rifampicin matched previous observations (Fig. 5d–h, Extended Data Fig. 5b). Menadione-treated WT cells exhibited a large subpopulation with Log(G/R) values near zero, consistent with growth arrest, alongside slowly replicating cells that progressively increased their Log(G/R) from 20 hours (Fig. 5d–h). Remarkably, WT cells treated with both menadione, and rifampicin exhibited growth-arrested bacteria, phenocopying rifampicin-treated *codY* mutant (Fig. 5d,h,l).

In the *codY* mutant, Log(G/R) patterns were consistent with previous observations (Fig. 5d,e–l, Extended Data Fig. 5b). Of note, in one replicate, a small subpopulation resumed growth—evidenced by increased Log(G/R) values—likely corresponding to a rifampicin-resistant mutant (Fig. 5j). Interestingly, the *codY* mutant treated with menadione alone was able to resume growth as early as 19 hours, suggesting a possible metabolic adjustment to elevated ROS. Taken together, these results indicate that *S. aureus*-derived ROS induced by rifampicin treatment result in persister formation, characterized by a growth-arrested phenotype.

## DISCUSSION

It is well established that *S. aureus* can survive antibiotic treatment by forming tolerant and persister cells [12]. Here, we uncover a previously unrecognized mechanism of persistence during rifampicin exposure, identifying CodY as a central regulator of oxidative stress metabolism and growth resumption. By engineering *S. aureus*-optimized fluorescent Timer-based reporters, we resolved at single-cell resolution the phenotypic heterogeneity underlying rifampicin response. We demonstrate that rifampicin exposure generates a heterogeneous population comprising resistant mutants, dying cells, and growth-arrested cells that fulfill the definition of persisters [10].

Timer and Timer^FAST^ reporters extend fluorescence-based approaches for studying bacterial recalcitrance. Timer^FAST^ carries the N42Q mutation, which accelerates chromophore maturation [28], thereby mitigating fluorescence dilution. Its faster maturation comes at the cost of reduced temporal resolution beyond three hours of growth arrest, whereas Timer is optimal for monitoring arrest over longer periods. We also introduced the V105A mutation into Timer^FAST^, which increases fluorescence quantum yield [27]. While this mutation could also enhance signal intensity of Timer, it would not address fluorescence dilution when bacterial division outpaces chromophore maturation. Our findings highlight the need of engineering Timer reporters with maturation kinetics tailored to pathogen biology and clinical contexts.

Combining Timer^FAST^ with live-cell microscopy enabled single-cell identification of rifampicin-induced persisters, which are otherwise masked in planktonic time-kill assays by the emergence of resistant mutants. We show that rifampicin treatment during exponential growth induces both resistant mutants and persisters, with the persister phenotype strongly enriched in the *codY* mutant. Growing evidences indicate that exogenous ROS and RNS can drive *S. aureus* toward antibiotic recalcitrance, including tolerance to rifampicin [14,35,36]. Antibiotics themselves are known to increase endogenous ROS level, thereby amplifying their killing effect beyond their direct targets [33,34,37]. Consistently, we observed robust ROS production during rifampicin exposure, which was markedly elevated in the *codY* mutant. CodY is a global transcriptional repressor responsive to GTP and branched-chain amino acids that fine-tunes metabolism, virulence, and stress responses [38–45]. Previous work has linked CodY to nitric oxide resistance in *S. aureus* [42] and to oxidative metabolism in *Streptococcus pneumoniae* [46]. Transcriptomic studies further show that *codY* mutation dysregulates catalase, peroxidases, and superoxide dismutase genes in a context-dependent manner [44,45], suggesting that multiple major regulators operating within a complex regulatory network. Collectively, these findings suggest that CodY protects against superoxide stress. In our study, failure of the *codY* mutant to restrain ROS under rifampicin pressure likely explains its reduced survival and pronounced persister phenotype. Despite the mutagenic potential of ROS [47], the *codY* mutant produced fewer resistant mutants, reflecting the predominance of non-replicating cells that cannot select mutations in the rifampicin resistance–determining region of the *rpoB* gene [48]. Menadione treatment confirmed that ROS detoxification is critical for recovery, as menadione-induced persisters resumed growth once treatment ends. Interestingly, CodY-deficient bacteria recovered more efficiently from menadione, suggesting either better elimination of menadione or a metabolic adaptation to high oxidative stress such as protein or DNA reparation [49]. Yet under rifampicin, these mutants generated almost exclusively persisters. Co-treatment of wild-type cells with rifampicin and menadione phenocopied this effect, suggesting that saturation of antioxidant defenses, combined with rifampicin’s target inhibition, shifts the balance entirely toward persistence. Similar threshold effects were observed with pyocyanin, a redox-active metabolite from *Pseudomonas aeruginosa*, which abolished growth regardless of CodY status [45].

In conclusion, our work establishes Timer and Timer^FAST^ as novel growth reporters for *S. aureus*, enabling high spatiotemporal resolution for studying recalcitrance. Using Timer^FAST^, we uncovered that CodY operates as a metabolic rheostat, fine-tuning the oxidative stress response during rifampicin exposure and thereby influencing the balance between resistance acquisition and persisters formation. Although antibiotic resistance complicates therapeutic decision-making, the majority of *S. aureus* clinical isolates remain susceptible to clinically approved antibiotics, including rifampicin. Recalcitrance emerge as an underappreciated contributor to treatment failure. Recent strategies including amino acid analogs [50] and antioxidants with *in vivo* efficacy [14] highlight the therapeutic potential of targeting persistence. By enabling precise single-cell analysis of growth heterogeneity, our Timer-based reporters open new possibilities for dissecting *S. aureus* recalcitrance in infection models and for developing strategies to eradicate persistent populations.

## MATERIALS AND METHODS

### Bacterial strains and growth conditions

*S. aureus* strains used in this study are listed in the Supplementary Table 1. For each experiment, strains were grown on tryptic soy agar (TSA) (22091-500; Sigma-Aldrich) or Mueller Hinton Agar (MHE) (413822; BioMérieux) plates and incubated for 24 hours at 37 °C. Liquid pre-cultures were prepared by inoculating a single colony into 2 ml of tryptic soy broth (TSB) (22092-500; Sigma-Aldrich) in 14 ml round-bottom culture tubes (352059; Falcon) and incubating overnight at 37°C with shaking at 225 rpm. To obtain exponential cultures, overnight cultures were diluted 1:200 into 15 ml of pre-warmed TSB in 50 ml conical tube (352070; Falcon) and incubated for 4.5 hours at 37 °C with shaking at 225 rpm. When appropriate, chloramphenicol (10368030; Fischer Scientific) was added at a final concentration of 20 µg/ml for plasmid selection. The minimum inhibitory concentration (MIC) of rifampicin was determined by agar diffusion using ETEST Rifampicin strips (412449; Biomérieux), following the manufacturer instructions. For treatment and selection of resistant mutants, culture media were supplemented with rifampicin (R3501-250MG; Sigma-Aldrich) at final concentration of 0.2 or 2 µg/ml, corresponding to 10x and 100x MIC, respectively.

### Rifampicin stock preparation and stability testing

Rifampicin powder (R3501-250MG; Sigma-Aldrich) was dissolved in DMSO (D12345, Fisher Scientific) to prepare a 10 mg/ml stock solution. Aliquots of 10 µl were snap-frozen and stored at -80°C. For each experimental replicate, a single aliquot was thawed at room temperature and diluted in sterile water to prepare a fresh working solution. The stability of stock solution was assessed weekly (see Supplementary Fig.1 for details).

### Molecular cloning of fluorescent reporters

Plasmids and primers used in this study are listed in Supplementary Tables 2 and 3. The Timer fluorescent reporter was constructed by site-directed mutagenesis of pSK265-DsRed [26]. All constructs were cloned into *S. aureus* RN4220 by electroporation, as previously described [26,51]. The S197T mutation was introduced by extension PCR using primers NP61 and NP62, and confirmed by DNA sequencing. This mutation resulted in plasmid pNP7, which enables constitutive expression of the Timer protein under the control of the *rpoB* promoter. The Timer^Fast^ fluorescent reporter was derived from pNP7 through two successive rounds of site-directed mutagenesis using primers NP74, NP75, NP76, and NP77. These steps resulted in plasmids pNP8 (carrying the S197T and N42Q mutations) and pNP9 (carrying S197T, N42Q, and V105A mutations). All mutations were verified by DNA sequencing, and full plasmid sequences were validated by outsourced long-read sequencing (Microsynth AG, Switzerland). Plasmids were subsequently introduced into *S. aureus* HG001 and SF8300 strains by electroporation [26,51].

### Agar plate imaging and analysis

A single colony of *S. aureus* expressing Timer, Timer^FAST^, DsRed or GFP were isolated on TSA plates and incubated for 48 hours at 37°C and brightfield images were acquired using a multimodal imaging system (iBright 1500FL; Fisher Scientific) at 24 and 48 hours of incubation. Fluorescence detection was performed using the Alexa Fluor 488 channel (50 ms exposure) for green emission and the Alexa Fluor 555 channel (100 ms exposure) for red emission. Brightfield images were captured using the visible light channel (400–800 nm) with a neutral density filter and 200 ms exposure. Images were acquired at 1.4x zoom and 1x1 resolution/sensitivity settings. Images were exported as TIFF-files and processed in Fiji software [52]. For each condition, three biological replicates were performed, and five colonies per plate were randomly analyzed. Within each colony, fluorescence intensity was measured for a single pixel in both green and red channels and corrected by subtracting background fluorescence from an adjacent area of the agar plate. Background-subtracted values were then used to calculate the fluorescence ratio as Log[(green MFI)/(red MFI)] (hereafter referred to as Log(G/R)) for Timer- and Timer^FAST^-expressing *S. aureus*.

### Live-cell imaging of *S. aureus* microcolony

The *S. aureus* microcolony model was adapted from [53]. Briefly, Hank’s Balanced Salt Solution (HBSS) (21-023-CM, Corning) and/or TSB were supplemented with 1% GTG agarose (LON50071; Lonza), boiled in a microwave, and maintained at 50°C in a dry bath to prevent solidification. Bacteria were harvested either in stationary phase (overnight cultures) or mid-exponential phase. Bacteria were washed twice with phosphate-buffered saline (PBS) (D8537-500ML; Sigma-Aldrich) and placed in an ultrasonic water bath for 1 minute to disperse cell aggregates. The bacterial suspension was then adjusted to an OD_600 nm_ of 0.5 (∼2.10^8^ cells/ml) and serially diluted to 10^-3^ in HBSS.

A 20 µL volume was deposited into each well of an 8-well chamber slide (µ-Slide 8 Well, 80826, Ibidi GmbH) and incubated at 37 °C for 10 minutes to allow the drop to dry. The bacteria were then embedded in agarose-supplemented medium and incubated at room temperature for 5 minutes to allow the agarose to solidify. Microcolony formation was monitored by time-lapse microscopy.

Live-cell imaging was performed on a spinning disk confocal microscope (Ti2 CSU-W1, Nikon) equipped with a 60× oil-immersion objective (CFI Plan Apo Lambda S 60×/1.40, MRD1605, Nikon) and a back-illuminated sCMOS camera (Prime BSI, Teledyne). Green and red fluorescence were excited with 488 nm (10 mW) and 561 nm (20 mW) lasers and collected through 525/50 nm and 600/52 nm bandpass filters, respectively, with 100 ms exposure. Brightfield images were acquired using differential interference contrast (DIC) with 50 ms exposure. Camera operated in CMS/12-bit mode for both DIC and fluorescence images. Environmental conditions were maintained with a stage-top incubator (OKO-Touch, Okolab, Pozzuoli, Italy) set to 37 °C with 60% relative humidity. Z-stacks were acquired every 20 min for up to 8 hours across five fields of view per well. Image acquisition was fully automated using NIS-Elements v5.30 (Nikon, France).

Images analysis was carried out using NIS-Elements software v5.30 (Nikon). Staphylococci were segmented using artificial intelligence—based models. Briefly, representative microcolonies were manually segmented from several images acquired in the green fluorescence channel, either from individual Z-stack planes (3D) or from maximum intensity projection images (2D). These manually annotated datasets were used to train distinct models using the Segment.ai module with 500 iterations and dynamic range correction applied. Prior to segmentation, images were pre-processed using Restore.ai, converted to 32-bit format, and segmented using the previously trained 2D or 3D Segment.ai models. For 2D analysis, maximum intensity projection was applied before pre-processing. Segmented microcolonies were assigned color-coded IDs and filtered based on circularity (range: 0.1–1.0) and size thresholds (2D: 0.5–500 µm²; 3D: 1.0–10,000 µm³). Objects touching the image border were excluded from the analysis. For each microcolony, area (2D) or volume (3D) was calculated from the binary layers. Mean fluorescence intensities (MFI) were extracted from the corresponding raw images. The fluorescence ratio for Timer or Timer^FAST^ was computed as Log(G/R) for each individual microcolony.

### Growth curves and generation time measurement

*S. aureus* strains expressing Timer and Timer^FAST^ were first grown overnight, then washed twice with PBS to remove residual rich medium. Cell aggregates were dispersed by sonication in an ultrasonic water bath for 1 minute. Following centrifugation, cells were resuspended in HBSS (21-023-CM; Corning) and adjusted to an absorbance of 0.5 at OD_600_ (∼2.10^8^ cells/ml). Ten microliters of the suspension were then inoculated into 90 µl of either TSB90 (9:1 TSB-HBSS) or TSB5 (5-95 TSB-HBSS) or pure HBSS in a flat bottom 96-well plate. The plate was incubated in multimodal plate reader (Infinite 200 PRO, Tecan) at 37°C with shaking at 225 rpm. Absorbance measurement at OD_600_ was recorded every 10 minutes. At 4-, 6- and 8-hours post-inoculation, one well per condition was collected, resuspend in 900 µl of PBS to be analyzed by flow cytometry as described hereafter. Generation times were calculated from growth curves using Prism software v10 (GraphPad software, Boston, MA). Absorbance values were converted into their natural logarithm ln(OD_600_) and plotted versus time (minutes). A linear regression was applied to the exponential phase. The slope of the resulting line (r), corresponding to the growth rate, was used to calculate the generation time (G) using the formula G=ln(2)/r.

### Flow cytometry experiment and analysis

For data collected in Fig. 1, *S. aureus* was cultured in a plate reader at 37°C and 225 RPM in TSB90 or TSB5 to the exponential phase (during 3 to 5 hours) and in HBSS overnight before sampling for cytometry. For data collected in Fig. 2 and Fig.3, *S. aureus* was cultured to the exponential phase and was then treated or not with rifampicin, sampling was done at time T0, T18 and T24 for cytometry.

For each sample, 2 µl of the suspension was resuspended in 1 ml of PBS and was sonicated 1 minute in an ultrasonic water bath to disperse cell aggregates prior to flow cytometry analysis. Bacteria were analyzed using a full spectrum flow cytometer (Northern Lights spectral flow cytometer, Cytek Biosciences). Approximately 30,000 events were acquired per sample at a flow rate of 10 µl/min. Bacterial populations were identified based on forward scatter (FSC) and side scatter (SSC) parameters. Fluorescence excitation was carried using lasers at 405 nm (100 mW), 488 nm (50 mW) and 640 nm (80 mW). Spectral detection was performed using Coarse Wavelength Division Multiplexing, capturing the emission spectrum in the 420-829 nm range. FSC and SSC parameters were set to selectively gate the *S. aureus* population, using a tube containing unstained *S. aureus* harboring the empty vector pNP4 (Supplementary Table 2) in PBS and PBS-only tube as negative control. Spectral overlap was corrected using single-stained reference controls. Reference spectra for DsRed, GFP and unstained cells were acquired using overnight culture of *S. aureus* strains. Live spectral unmixing was performed using the SpectroFlo software CS Version 1.4.1 (Cytek Biosciences). The unmixing algorithm assigned fluorescence signals to their fluorophores based on the acquired reference spectra minimizing spectral bleed-through. Gating strategies were defined using the fluorescence-minus-one and unstained controls to ensure accurate and specific detection of Timer and Timer^FAST^ fluorescence. Data were analyzed using FlowJo software v10.10.0 (Becton Dickinson). Fluorescence intensity values were used to extract either scatter plots of green fluorescence as a function of red fluorescence or frequency distribution of the log-transformed fluorescence ratio and Log(G/R).

To distinguish subpopulations based on physiological state during rifampicin treatment, additional controls were included. Cultures of *S. aureus* expressing Timer^FAST^ in either mid-exponential (∼4h30) or stationary phases (∼24h) were used to define two primary populations: Q1, corresponding to actively growing cells (high green, low red fluorescence), and Q2, corresponding to non-growing cells (high green, high red fluorescence), based on their respective fluorescence intensity profiles. A heat-killed control, generated by diluting an overnight culture to an OD_600_ of 0.5 and treated at 100°C for 5 minutes, was included as a non-viable, autofluorescence-free reference to define Q3 population (see Supplementary Fig.2 for details). Additionally, 2µg/ml propidium iodide (PI) (P3566; Fisher Scientific) was added to identify permeabilized cells. Spectral unmixing for PI was performed using heat-killed, PI-stained cells treated with PI as a positive control.

### Killing assay

Bacteria were grown in TSB to exponential phase as described above. Bacterial load at baseline was measured on TSA using an automatic diluter and plater (EasySpiral dilute, Interscience, France) and an automatic colony counter (Scan 4000, Interscience). A volume of 2 ml of exponential culture was mock-treated with 10 µl of sterile water containing 0.5% DMSO or treated with 10 µl of rifampicin (R3501-250MG; Sigma-Aldrich) to reach 10x MIC (0.2 μg/ml) or 100x MIC (2 µg/ml), and/or 100 µM menadione (M5625-25G; Sigma-Aldrich) during 18 hours or more as indicated. At the indicated time points, a 500 µL aliquot were collected, washed once with 500 µl of PBS and processed for either bacterial load quantification, microcolony imaging or flow cytometry analysis. CFU quantification on agar plate was performed using an automatic colony counter (Scan 4000, Interscience). All experiments were performed in biological triplicates or more. Reported averages and standard deviations are representative of three or more independent biological replicates.

### Resistance rate assay

Bacteria were grown in TSB to exponential phase (∼2.10^8^ cells/mL) as described above. Cultures were then concentrated ten times by centrifugation at 4,000g during 4 min and resuspension in TSB to reach ∼2.10^9^ cells/mL. 200 µL of this suspension was then plated on TSA and TSA supplemented with 2 µg/ml of rifampicin (TSA^RIF^) using an automatic diluter and plater (EasySpiral dilute, Interscience, France) and the next day bacterial load at baseline was measured using a colony counter (Scan 4000, Interscience). The resistance rate was calculated by dividing the CFU count on TSA^RIF^ by the CFU count on TSA. Three independent biological replicates were performed on separate days.

### Reactive oxygen species (ROS) level measurement

ROS level was assessed using the oxidation-sensitive fluorescent probe 2,7-dichlorofluorescin diacetate (H_2_DCFDA) (D6883-50MG; Sigma-Aldrich) [54]. Bacteria were grown in TSB to exponential or stationary phase as described above. H_2_DCFDA was added to the culture at a final concentration of 20 µM, with 5% DMSO, rifampicin at 2 µg/ml and/or menadione at 100 µM as indicated. After an incubation period of 30 minute at 37°C, fluorescence was measured using a multimodal plate reader (Infinite 200 Pro, Tecan) with excitation at 488 nm and emission at 530 nm. For quantification of ROS level at H18, H_2_DCFDA was added at H0 and signal was recorded every hour for 18 hours.

### Statistical analysis

Statistical methods and sample sizes (n) are detailed in the figure legends for each experiment. Statistical analyses and data visualization were performed using Prism v10 (GraphPad software) or FlowJo v10.10.0 (Becton Dickinson). For microscopy-based analysis, *n* refers to individual bacteria or microcolonies, with data derived from at least three independent experiments. For flow cytometry and bacterial load quantification, *n* represents the number of biological replicates. In plate reader assays, *n* corresponds to individual wells within a 96-well plate, with all experiments conducted in a minimum of three independent replicates. Statistical significance was defined as *p* < 0.05. Error bars represent either the standard deviation (SD) of the mean or the interquartile range (IQR) of the median, as indicated in the figure legend.

---

**Extended data Fig. 1.**
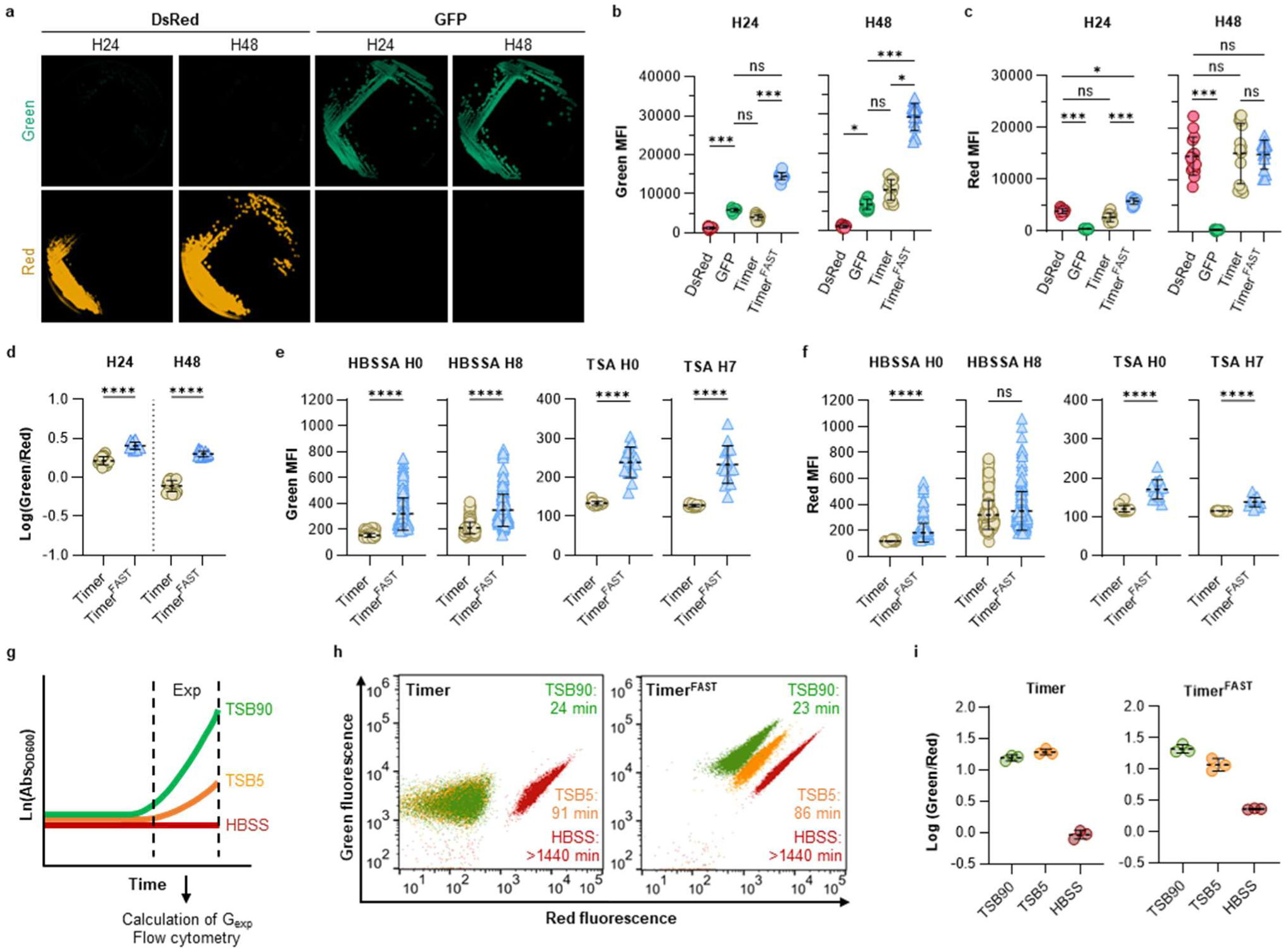
Spectral dynamics of *S. aureus* expressing GFP, DsRed, Timer and Timer^FAST^. **a**, Colonies of *S. aureus* SF8300 constitutively expressing GFP or DsRed grown for 24 and 48 hours on TSA plates supplemented with chloramphenicol. **b–d**, Mean fluorescence intensity (MFI) in the green (b) and red (c) channels, or fluorescence ratio (d) from single colonies illustrated in Fig. 1a and panel (a). Each dot represents a single colony acquired across three independent biological experiments. Dotted lines and error bars: mean ± SD. Mann–Whitney U test: \**P* < 0.05; \*\*\**P* < 0.001; \*\*\*\**P* < 0.0001; ns, not significant. **e–f**, Green (e) and red (f) MFI of Timer- and Timer^FAST^-expressing *S. aureus* at 0 or 7 hours, corresponding to experiments illustrated in Fig. 1c,d. Dotted lines and errors bars: mean ± SD. Mann-Whitney U test: \*\*\*\**P* < 0.0001; ns, not significant. **g**, Workflow for flow cytometry experiments during exponential growth. Bacteria were grown in TSB90 (9:1 TSB:HBSS), TSB5 (5:95 TSB:HBSS), or HBSS in a multimodal plate reader. Generation time during exponential phase (G_exp_) was calculated from Ln(Abs_OD600_). **h**, Flow cytometry analysis of green and red MFI of Timer- and Timer^FAST^-expressing *S. aureus* grown in TSB90, TSB5, and HBSS. G_exp_ values are indicated for each condition. Data are from one representative experiment out of three independent biological replicates. **i**, Fluorescence ratios of Timer- and Timer^FAST^-expressing *S. aureus* measured by flow cytometry, illustrated in (h). Dotted lines and error bars: mean ± SD. Data correspond to three independent biological replicates (n = 21,480 events for TSB90; n = 23,625 events for TSB5; n = 29,201 events for HBSS). HBSS: Hank’s balanced salt solution. HBSSA: Hank’s balanced salt agar. TSA: tryptic soy agar. TSB: tryptic soy broth.

**Extended data Fig. 2.**
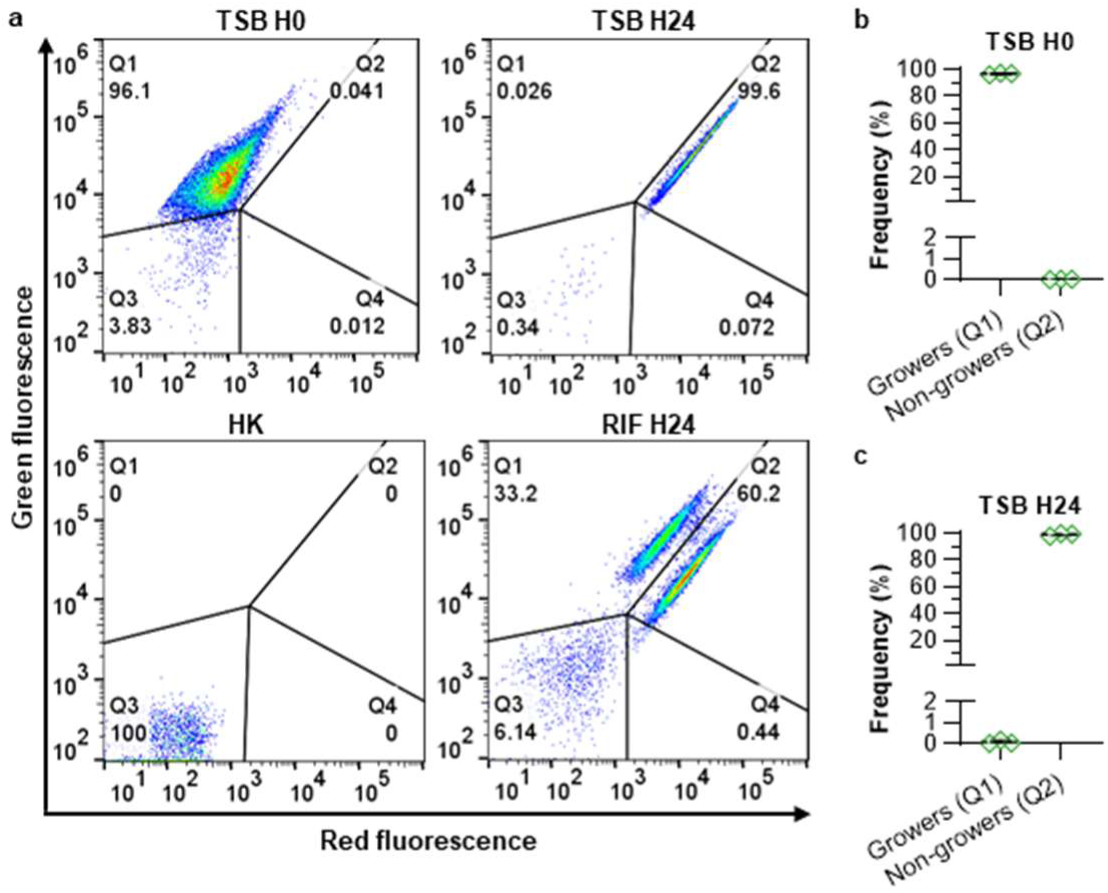
Flow cytometry gating strategy of Timer^FAST^-expressing *S. aureus*. **a**, Flow cytometry analysis of Timer^FAST^-expressing *S. aureus* grown in exponential phase (TSB H0), in stationary phase (TSB H24), treated 24 hours with 10x MIC rifampicin (RIF H24), or heat killed (HK). Q1: growers. Q2: non-growers. Q3: dead cells. Q4: none. Data shown are from one representative of three independent flow cytometry experiments. **b-c**, Frequency of growers (Q1) and non-growers (Q2) after 4.5 (b) or 28.5 (c) hours of incubation in TSB using the gating strategy is shown in (a). Each dot represents an independent experiment. Dotted lines and errors bars: mean ± SD.

**Extended Data Fig. 3.**
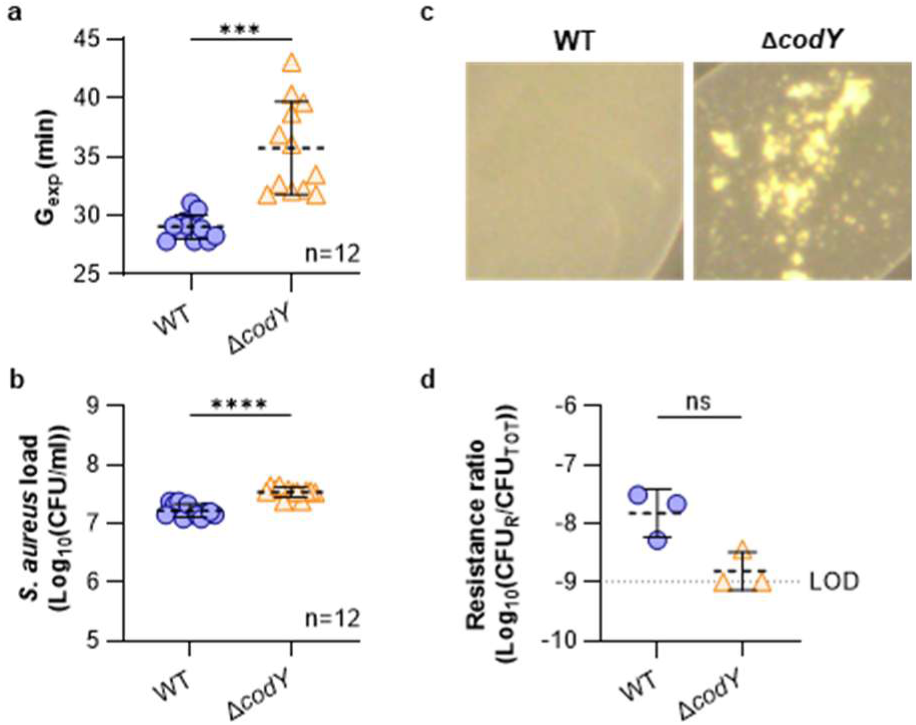
Growth behaviour of *S. aureus* wild-type (WT) and *codY* mutant strains. **a**, Generation times during exponential growth (G_exp_) of *S. aureus* HG001 WT and *codY* mutant strains grown in TSB. Each dot represents a value calculated from OD measurements acquired from a single well across three independent biological experiments. Dotted lines and errors bars: mean ± SD. Mann-Whitney U test: *** P < 0.001. b, *S. aureus* load for HG001 WT and *codY* mutant strains grown for 4.5 hours in TSB (corresponding to the H0 time point in Fig. 3a). Each dot represents the value of a single broth from six independent experiments. Dotted lines and error bars: mean ± SD. Mann-Whitney U test: **** P < 0.001. **c**, Representative illustration of macroscopic aggregates observed in overnight cultures of the *S. aureus* HG001 *codY* mutant strain compared with its isogenic WT strain. **d**, Proportion of rifampicin-resistant CFU (CFU_R_) relative to total CFU (CFU_TOT_) for *S. aureus* HG001 WT and *codY* mutant strains. Each dot represents a single value from three independent experiments. LOD: limit of detection. Dotted lines and error bars: mean ± SD. Mann-Whitney U test: ns, not significant.

**Extended Data Fig. 4.**
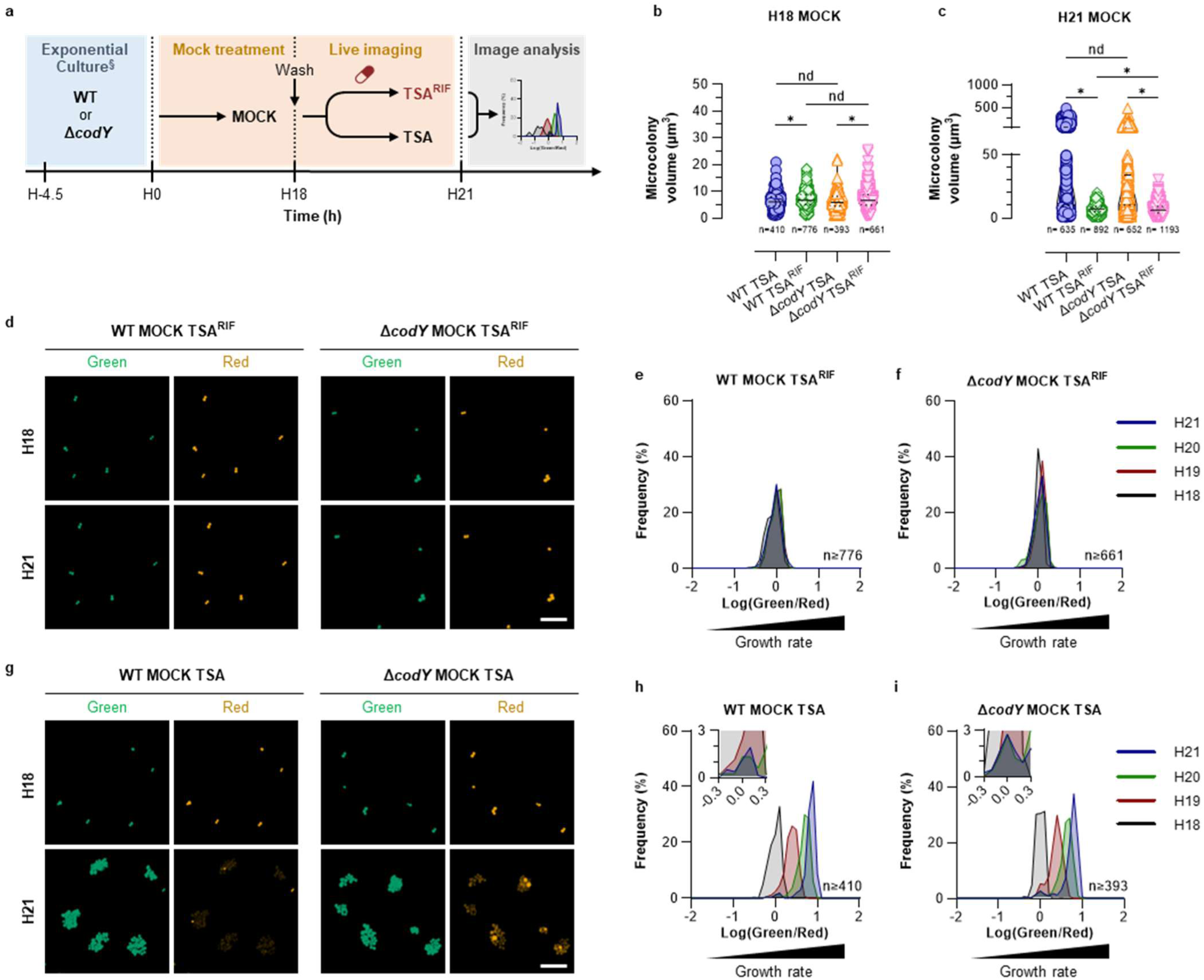
*codY* deficiency increase the non-growing population during rifampicin treatment. **a**, Experimental workflow of Timer^FAST^-expressing *S. aureus* HG001 wild-type (WT) and *codY* mutant strains treated with DMSO (MOCK) in TSB during 18 hours (H18), embedded in TSA or in TSA supplemented with 100x minimum inhibitory concentration rifampicin (TSA^RIF^), and analyzed by live-cell confocal microscopy (b-i). ^§^Exponential cultures were prepared as described in Fig. 3a. **b,c**, *S. aureus* microcolony volume at 18 (H18, b) and 21 (H21, c) hours. Values (n) indicate the number of microcolony measured across three independent biological experiments. Solid and dotted lines: median ± IQR. Kruskal-Wallis test with Benjamini and Hochberg false discovery rate correction for multiple comparisons: * Q < 0.05; nd, not discovered. **d,g**, Representative images of *S. aureus* microcolony embedded in TSA^RIF^ (d), or TSA (g), at 18 (H18) and 21 (H21) hours. Green channel: Ex 488 nm, Em 525/50 nm. Red channel: Ex 561 nm, Em 600/52 nm. Scale bar: 10 µm. e,f,h,i, Green to red fluorescence ratios of *S. aureus* microcolonies from 18 (H18) to 21 (H21) hours. The number of microcolonies (n) analyzed across three independent biological experiments is indicated on the graph. Insets depict an enlarged view of the region of the x-axis near zero.

**Extended Data Fig. 5.**
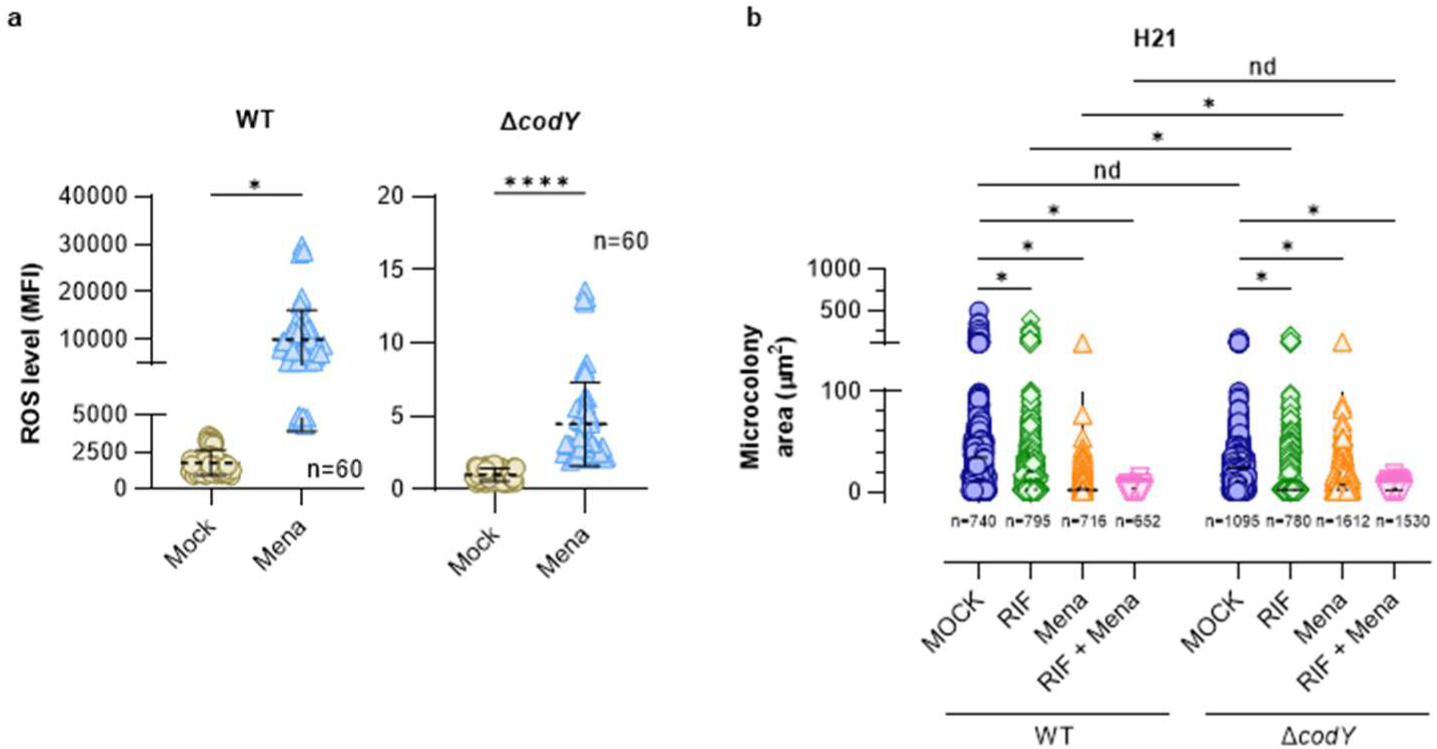
Reactive oxygen species promotes a non-growing phenotype of *S. aureus* during rifampicin treatment. **a**, Reactive oxygen species (ROS) level in *S. aureus* HG001 wild-type (WT) and *codY* mutant strains treated with DMSO (MOCK) or 100 µM menadione (Mena). Each dot represents a single well acquired across three independent biological experiments. Dotted lines and error bars: mean ± SD. Mann-Whitney U test: **** P < 0.0001; ns, not significant. **b**, *S. aureus* microcolony area at 21 hours (H21). Values (n) indicate the number of microcolony measured across three independent biological experiments. Dotted lines and error bars: median ± IQR. Kruskal-Wallis test with Benjamini and Hochberg false discovery rate correction for multiple comparisons; *Q < 0.05; nd, not discovered.

## DATA AVAILABILITY

The data support the findings of this study are available in the Source Data file.

## ACKNOWLEDGMENTS

The *Staphylococcus aureus* HG001 strain and its derivatives mutants were kindly provided by Prof. Christiane Wolz (University of Tübingen, Germany). The authors gratefully acknowledge Alison Carteron and Inès Verneuil for their technical assistance. This work was supported by the “Fondation Université Jean Monnet” as part of the YAPSTAPH project (APP2023).

## AUTHOR CONTRIBUTIONS

Conceptualization: NPO, KR, JG, NPE, EBN, POV; Methodology: NPO, KR, JG, NPE, POV; Investigation: NPO (cloning, time-kill assay, live-cell confocal microscopy, flow cytometry), JG (time-kill assay, flow cytometry); Formal analysis: NPO, POV; Visualization: NPO, POV; Writing – original draft: NPO, POV; Writing – review & editing: JG, NPE, EBN; Supervision: JG, EBN, POV; Funding acquisition: POV

## REFERENCES

[1] GBD 2019 Antimicrobial Resistance Collaborators. Global mortality associated with 33 bacterial pathogens in 2019: a systematic analysis for the Global Burden of Disease Study 2019. Lancet 2022:S0140-6736(22)02185-7. 10.1016/S0140-6736(22)02185-7.

[2] Tong SYC, Davis JS, Eichenberger E, Holland TL, Fowler VG. Staphylococcus aureus infections: epidemiology, pathophysiology, clinical manifestations, and management. Clin Microbiol Rev 2015;28:603–61. 10.1128/CMR.00134-14.

[3] Verhoeven PO, Gagnaire J, Botelho-Nevers E, Grattard F, Carricajo A, Lucht F, Pozzetto B, Berthelot P. Detection and clinical relevance of Staphylococcus aureus nasal carriage: an update. Expert Rev Anti Infect Ther 2014;12:75–89. 10.1586/14787210.2014.859985.

[4] Nicolas R, Carricajo A, Morel J, Rigaill J, Grattard F, Guezzou S, Audoux E, Campisi S, Favre J-P, Berthelot P, Verhoeven PO, Botelho-Nevers E. Evaluation of effectiveness and compliance with the mupirocin nasal ointment part of Staphylococcus aureus decolonization in real life using UPLC-MS/MS mupirocin quantification. J Antimicrob Chemother 2020;75:1623–30. 10.1093/jac/dkaa025.

[5] Tacconelli E, Carrara E, Savoldi A, Harbarth S, Mendelson M, Monnet DL, Pulcini C, Kahlmeter G, Kluytmans J, Carmeli Y, Ouellette M, Outterson K, Patel J, Cavaleri M, Cox EM, Houchens CR, Grayson ML, Hansen P, Singh N, Theuretzbacher U, Magrini N, WHO Pathogens Priority List Working Group. Discovery, research, and development of new antibiotics: the WHO priority list of antibiotic-resistant bacteria and tuberculosis. Lancet Infect Dis 2018;18:318–27. 10.1016/S1473-3099(17)30753-3.

[6] Huemer M, Mairpady Shambat S, Brugger SD, Zinkernagel AS. Antibiotic resistance and persistence-Implications for human health and treatment perspectives. EMBO Rep 2020;21:e51034. 10.15252/embr.202051034.

[7] Conlon BP, Nakayasu ES, Fleck LE, LaFleur MD, Isabella VM, Coleman K, Leonard SN, Smith RD, Adkins JN, Lewis K. Activated ClpP kills persisters and eradicates a chronic biofilm infection. Nature 2013;503:365–70. 10.1038/nature12790.

[8] Conlon BP, Rowe SE, Gandt AB, Nuxoll AS, Donegan NP, Zalis EA, Clair G, Adkins JN, Cheung AL, Lewis K. Persister formation in Staphylococcus aureus is associated with ATP depletion. Nat Microbiol 2016;1:16051. 10.1038/nmicrobiol.2016.51.

[9] Huemer M, Mairpady Shambat S, Bergada-Pijuan J, Söderholm S, Boumasmoud M, Vulin C, Gómez-Mejia A, Antelo Varela M, Tripathi V, Götschi S, Marques Maggio E, Hasse B, Brugger SD, Bumann D, Schuepbach RA, Zinkernagel AS. Molecular reprogramming and phenotype switching in Staphylococcus aureus lead to high antibiotic persistence and affect therapy success. Proc Natl Acad Sci U S A 2021;118:e2014920118. 10.1073/pnas.2014920118.

[10] Balaban NQ, Helaine S, Lewis K, Ackermann M, Aldridge B, Andersson DI, Brynildsen MP, Bumann D, Camilli A, Collins JJ, Dehio C, Fortune S, Ghigo J-M, Hardt W-D, Harms A, Heinemann M, Hung DT, Jenal U, Levin BR, Michiels J, Storz G, Tan M-W, Tenson T, Van Melderen L, Zinkernagel A. Definitions and guidelines for research on antibiotic persistence. Nat Rev Microbiol 2019;17:441–8. 10.1038/s41579-019-0196-3.

[11] Bakkeren E, Diard M, Hardt W-D. Evolutionary causes and consequences of bacterial antibiotic persistence. Nat Rev Microbiol 2020;18:479–90. 10.1038/s41579-020-0378-z.

[12] Helaine S, Conlon BP, Davis KM, Russell DG. Host stress drives tolerance and persistence: The bane of anti-microbial therapeutics. Cell Host Microbe 2024;32:852–62. 10.1016/j.chom.2024.04.019.

[13] Peyrusson F, Varet H, Nguyen TK, Legendre R, Sismeiro O, Coppée J-Y, Wolz C, Tenson T, Van Bambeke F. Intracellular Staphylococcus aureus persisters upon antibiotic exposure. Nat Commun 2020;11:2200. 10.1038/s41467-020-15966-7.

[14] Rowe SE, Wagner NJ, Li L, Beam JE, Wilkinson AD, Radlinski LC, Zhang Q, Miao EA, Conlon BP. Reactive oxygen species induce antibiotic tolerance during systemic Staphylococcus aureus infection. Nat Microbiol 2020;5:282–90. 10.1038/s41564-019-0627-y.

[15] Beam JE, Wagner NJ, Lu K-Y, Parsons JB, Fowler VG, Rowe SE, Conlon BP. Inflammasome-mediated glucose limitation induces antibiotic tolerance in Staphylococcus aureus. iScience 2023;26:107942. 10.1016/j.isci.2023.107942.

[16] Leimer N, Rachmühl C, Palheiros Marques M, Bahlmann AS, Furrer A, Eichenseher F, Seidl K, Matt U, Loessner MJ, Schuepbach RA, Zinkernagel AS. Nonstable Staphylococcus aureus Small-Colony Variants Are Induced by Low pH and Sensitized to Antimicrobial Therapy by Phagolysosomal Alkalinization. J Infect Dis 2016;213:305–13. 10.1093/infdis/jiv388.

[17] Vulin C, Leimer N, Huemer M, Ackermann M, Zinkernagel AS. Prolonged bacterial lag time results in small colony variants that represent a sub-population of persisters. Nat Commun 2018;9:4074. 10.1038/s41467-018-06527-0.

[18] Häffner N, Bär J, Dengler Haunreiter V, Mairpady Shambat S, Seidl K, Crosby HA, Horswill AR, Zinkernagel AS. Intracellular Environment and agr System Affect Colony Size Heterogeneity of Staphylococcus aureus. Front Microbiol 2020;11:1415. 10.3389/fmicb.2020.01415.

[19] Marro FC, Laurent F, Josse J, Blocker AJ. Methods to monitor bacterial growth and replicative rates at the single-cell level. FEMS Microbiol Rev 2022;46:fuac030. 10.1093/femsre/fuac030.

[20] Claudi B, Spröte P, Chirkova A, Personnic N, Zankl J, Schürmann N, Schmidt A, Bumann D. Phenotypic variation of Salmonella in host tissues delays eradication by antimicrobial chemotherapy. Cell 2014;158:722–33. 10.1016/j.cell.2014.06.045.

[21] Personnic N, Striednig B, Lezan E, Manske C, Welin A, Schmidt A, Hilbi H. Quorum sensing modulates the formation of virulent Legionella persisters within infected cells. Nat Commun 2019;10:5216. 10.1038/s41467-019-13021-8.

[22] Personnic N, Striednig B, Hilbi H. Quorum sensing controls persistence, resuscitation, and virulence of Legionella subpopulations in biofilms. ISME J 2021;15:196–210. 10.1038/s41396-020-00774-0.

[23] Lang JC, Seiß EA, Moldovan A, Müsken M, Sauerwein T, Fraunholz M, Müller AJ, Goldmann O, Medina E. A Photoconvertible Reporter System for Bacterial Metabolic Activity Reveals That Staphylococcus aureus Enters a Dormant-Like State to Persist within Macrophages. mBio 2022;13:e0231622. 10.1128/mbio.02316-22.

[24] Matz MV, Fradkov AF, Labas YA, Savitsky AP, Zaraisky AG, Markelov ML, Lukyanov SA. Fluorescent proteins from nonbioluminescent Anthozoa species. Nat Biotechnol 1999;17:969–73. 10.1038/13657.

[25] Baird GS, Zacharias DA, Tsien RY. Biochemistry, mutagenesis, and oligomerization of DsRed, a red fluorescent protein from coral. Proc Natl Acad Sci U S A 2000;97:11984–9. 10.1073/pnas.97.22.11984.

[26] Caire R, Audoux E, Thomas M, Dalix E, Peyron A, Rodriguez K, Pordone N, Guillemot J, Dickerscheit Y, Marotte H, Vandenesch F, Laurent F, Josse J, Verhoeven PO. YAP promotes cell-autonomous immune responses to tackle intracellular Staphylococcus aureus in vitro. Nat Commun 2022;13:6995. 10.1038/s41467-022-34432-0.

[27] Terskikh A, Fradkov A, Ermakova G, Zaraisky A, Tan P, Kajava AV, Zhao X, Lukyanov S, Matz M, Kim S, Weissman I, Siebert P. “Fluorescent Timer”: Protein That Changes Color with Time. Science 2000;290:1585–8. 10.1126/science.290.5496.1585.

[28] Bevis BJ, Glick BS. Rapidly maturing variants of the Discosoma red fluorescent protein (DsRed). Nat Biotechnol 2002;20:83–7. 10.1038/nbt0102-83.

[29] Wang C, Fang R, Zhou B, Tian X, Zhang X, Zheng X, Zhang S, Dong G, Cao J, Zhou T. Evolution of resistance mechanisms and biological characteristics of rifampicin-resistant Staphylococcus aureus strains selected in vitro. BMC Microbiol 2019;19:220. 10.1186/s12866-019-1573-9.

[30] Lester W. Rifampin: a semisynthetic derivative of rifamycin--a prototype for the future. Annu Rev Microbiol 1972;26:85–102. 10.1146/annurev.mi.26.100172.000505.

[31] Zhu J-H, Wang B-W, Pan M, Zeng Y-N, Rego H, Javid B. Rifampicin can induce antibiotic tolerance in mycobacteria via paradoxical changes in rpoB transcription. Nat Commun 2018;9:4218. 10.1038/s41467-018-06667-3.

[32] Mlynek KD, Bulock LL, Stone CJ, Curran LJ, Sadykov MR, Bayles KW, Brinsmade SR. Genetic and Biochemical Analysis of CodY-Mediated Cell Aggregation in Staphylococcus aureus Reveals an Interaction between Extracellular DNA and Polysaccharide in the Extracellular Matrix. J Bacteriol 2020;202:e00593–19. 10.1128/JB.00593-19.

[33] Zhang J, Xu J, Lei H, Liang H, Li X, Li B. The development of variation-based rifampicin resistance in Staphylococcus aureus deciphered through genomic and transcriptomic study. J Hazard Mater 2023;442:130112. 10.1016/j.jhazmat.2022.130112.

[34] Dwyer DJ, Belenky PA, Yang JH, MacDonald IC, Martell JD, Takahashi N, Chan CTY, Lobritz MA, Braff D, Schwarz EG, Ye JD, Pati M, Vercruysse M, Ralifo PS, Allison KR, Khalil AS, Ting AY, Walker GC, Collins JJ. Antibiotics induce redox-related physiological alterations as part of their lethality. Proc Natl Acad Sci U S A 2014;111:E2100–2109. 10.1073/pnas.1401876111.

[35] Beam JE, Wagner NJ, Shook JC, Bahnson ESM, Fowler VG, Rowe SE, Conlon BP. Macrophage-Produced Peroxynitrite Induces Antibiotic Tolerance and Supersedes Intrinsic Mechanisms of Persister Formation. Infect Immun 2021;89:e0028621. 10.1128/IAI.00286-21.

[36] Peyrusson F, Nguyen TK, Najdovski T, Van Bambeke F. Host Cell Oxidative Stress Induces Dormant Staphylococcus aureus Persisters. Microbiol Spectr 2022;10:e0231321. 10.1128/spectrum.02313-21.

[37] Kohanski MA, Dwyer DJ, Hayete B, Lawrence CA, Collins JJ. A common mechanism of cellular death induced by bactericidal antibiotics. Cell 2007;130:797–810. 10.1016/j.cell.2007.06.049.

[38] Pohl K, Francois P, Stenz L, Schlink F, Geiger T, Herbert S, Goerke C, Schrenzel J, Wolz C. CodY in Staphylococcus aureus: a regulatory link between metabolism and virulence gene expression. J Bacteriol 2009;191:2953–63. 10.1128/JB.01492-08.

[39] Majerczyk CD, Dunman PM, Luong TT, Lee CY, Sadykov MR, Somerville GA, Bodi K, Sonenshein AL. Direct targets of CodY in Staphylococcus aureus. J Bacteriol 2010;192:2861–77. 10.1128/JB.00220-10.

[40] Geiger T, Francois P, Liebeke M, Fraunholz M, Goerke C, Krismer B, Schrenzel J, Lalk M, Wolz C. The stringent response of Staphylococcus aureus and its impact on survival after phagocytosis through the induction of intracellular PSMs expression. PLoS Pathog 2012;8:e1003016. 10.1371/journal.ppat.1003016.

[41] Corrigan RM, Bellows LE, Wood A, Gründling A. ppGpp negatively impacts ribosome assembly affecting growth and antimicrobial tolerance in Gram-positive bacteria. Proc Natl Acad Sci U S A 2016;113:E1710–1719. 10.1073/pnas.1522179113.

[42] Grosser MR, Weiss A, Shaw LN, Richardson AR. Regulatory Requirements for Staphylococcus aureus Nitric Oxide Resistance. J Bacteriol 2016;198:2043–55. 10.1128/JB.00229-16.

[43] Brinsmade SR. CodY, a master integrator of metabolism and virulence in Gram-positive bacteria. Curr Genet 2017;63:417–25. 10.1007/s00294-016-0656-5.

[44] Gao Y, Poudel S, Seif Y, Shen Z, Palsson BO. Elucidating the CodY regulon in Staphylococcus aureus USA300 substrains TCH1516 and LAC. mSystems 2023;8:e0027923. 10.1128/msystems.00279-23.

[45] Martini AM, Alexander SA, Khare A. Mutations in the Staphylococcus aureus Global Regulator CodY confer tolerance to an interspecies redox-active antimicrobial. PLoS Genet 2025;21:e1011610. 10.1371/journal.pgen.1011610.

[46] Hajaj B, Yesilkaya H, Shafeeq S, Zhi X, Benisty R, Tchalah S, Kuipers OP, Porat N. CodY Regulates Thiol Peroxidase Expression as Part of the Pneumococcal Defense Mechanism against H2O2 Stress. Front Cell Infect Microbiol 2017;7:210. 10.3389/fcimb.2017.00210.

[47] Carvajal-Garcia J, Samadpour AN, Hernandez Viera AJ, Merrikh H. Oxidative stress drives mutagenesis through transcription-coupled repair in bacteria. Proc Natl Acad Sci U S A 2023;120:e2300761120. 10.1073/pnas.2300761120.

[48] Guérillot R, Gonçalves da Silva A, Monk I, Giulieri S, Tomita T, Alison E, Porter J, Pidot S, Gao W, Peleg AY, Seemann T, Stinear TP, Howden BP. Convergent Evolution Driven by Rifampin Exacerbates the Global Burden of Drug-Resistant Staphylococcus aureus. mSphere 2018;3:e00550–17. 10.1128/mSphere.00550-17.

[49] Ezraty B, Gennaris A, Barras F, Collet J-F. Oxidative stress, protein damage and repair in bacteria. Nat Rev Microbiol 2017;15:385–96. 10.1038/nrmicro.2017.26.

[50] Huang D, Kang X, Yin Z, Zhao D, Ning Y, Liu H, Li F, Xie W, Li G, Wang X. On-Site Serine Delivery Drives Fermentation Pathway Reprogramming to Reverse Intracellular Staphylococcus aureus Persistence. ACS Nano 2025. 10.1021/acsnano.5c06864.

[51] Monk IR, Shah IM, Xu M, Tan M-W, Foster TJ. Transforming the untransformable: application of direct transformation to manipulate genetically Staphylococcus aureus and Staphylococcus epidermidis. mBio 2012;3:e00277–11. 10.1128/mBio.00277-11.

[52] Schindelin J, Arganda-Carreras I, Frise E, Kaynig V, Longair M, Pietzsch T, Preibisch S, Rueden C, Saalfeld S, Schmid B, Tinevez J-Y, White DJ, Hartenstein V, Eliceiri K, Tomancak P, Cardona A. Fiji: an open-source platform for biological-image analysis. Nat Methods 2012;9:676–82. 10.1038/nmeth.2019.

[53] Personnic N, Striednig B, Hilbi H. Single Cell Analysis of Legionella and Legionella-Infected Acanthamoeba by Agarose Embedment. Methods Mol Biol 2019;1921:191–204. 10.1007/978-1-4939-9048-1_12.

[54] Wang H, Joseph JA. Quantifying cellular oxidative stress by dichlorofluorescein assay using microplate reader. Free Radic Biol Med 1999;27:612–6. 10.1016/s0891-5849(99)00107-0.

