## Supplementary Information for "Rifampicin-induced *Staphylococcus aureus* persister formation is driven by CodY regulon and oxidative stress level"

SUPPLEMENTARY FIGURES 1-2

SUPPLEMENTARY TABLES 1-3

REFERENCES

SUPPLEMENTARY FIGURES

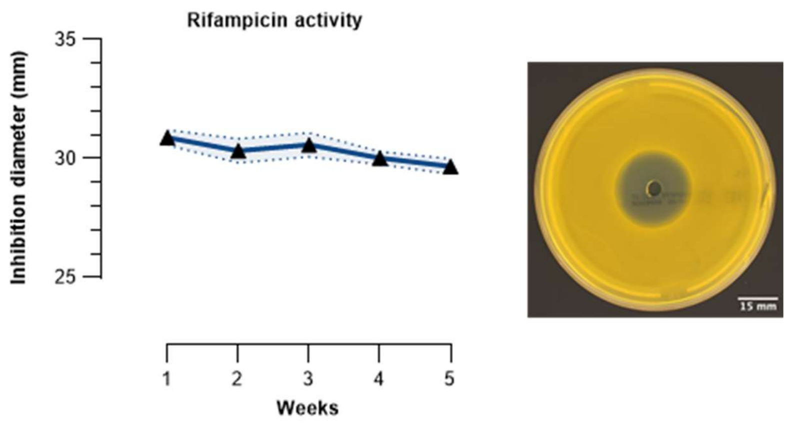


Supplementary Fig. 1 | Rifampicin stock preparation and stability testing

To assess the stability of the rifampicin stock solution, *S. aureus* strains were grown overnight in liquid culture and adjusted to an OD600 of 0.5 (~2 × $10^8$ CFU/ml). Mueller Hinton agar (MHE) plates were surface-flooded with 2 ml of the bacterial suspension, and excess liquid was removed. A 5 mm well was punched at the center of each plate, into which 50 µl of the rifampicin working solution was dispensed. Plates were incubated overnight at 37 °C. Zones of inhibition were imaged using a plate imaging system (Scan 4000, Interscience), and inhibition diameters were measured with Fiji software (https://fiji.sc/#). Stability testing was performed weekly over a five-week period, with each measurement done in triplicate. Rifampicin retained its antibacterial activity throughout this period under the described storage conditions (see results below). Scale bar: 15 mm.

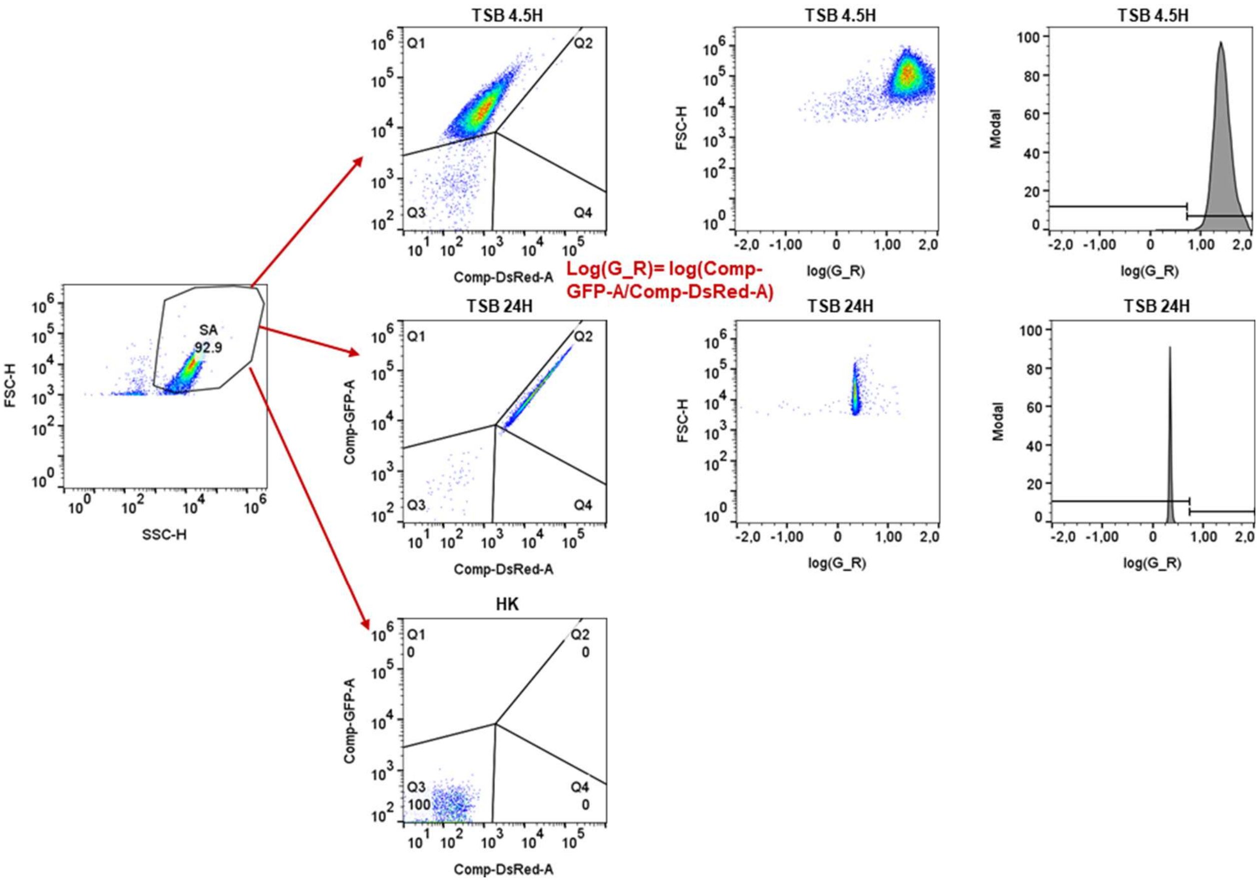


Supplementary Fig. 2 | Flow cytometry experiment and analysis

Gating was defined as shown in the figure. TSB 4.5 h: growing bacteria (high green, low red fluorescence) mapped to Q1. TSB 24 h: non-growing bacteria (high green, high red fluorescence) mapped to Q2. HK (heat-killed): autofluorescence-free bacteria used as a reference to define the Q3 population.

SUPPLEMENTARY TABLES

Supplementary Table 1 | Bacterial strains used in this study.

| **Strain** | **Characteristics** | **Reference** |
|---|---|---|
| RN4220 | *S. aureus* restriction deficient cloning host NCTC8325 | [1] |
| HG001 WT | HG001*, S. aureus* NCTC8325 *rsbU* repaired | [2] |
| HG001 Δ*codY::tetM* | HG001 *codY::tetM* mutant | [2] |
| SF8300 WT | SF8300*, S. aureus* minimally passaged clinical isolate belonging to USA300 clone | [3] |
| pSTAH54 | SF8300 expressing Timer under *rpoB* promotor, chloramphenicol resistance (*cat*) | This study |
| STAL44 | SF8300 expressing Timer$^{FAST}$ under *rpoB* promotor, chloramphenicol resistance (*cat*) | This study |
| STAJ42 | SF8300 containing the empty vector pNP4 | This study |
| STAG48 | SF8300 expressing DsRed under *rpoB* promotor, chloramphenicol resistance (*cat*) | This study |
| STAK01 | SF8300 expressing GFP under *rpoB* promotor, chloramphenicol resistance (*cat*) | This study |
| STAO75 | HG001 WT expressing Timer$^{FAST}$ under *rpoB* promotor, chloramphenicol resistance (*cat*) | This study |
| STAP42 | HG001 Δ*codY* expressing Timer$^{FAST}$ under *rpoB* promotor, chloramphenicol resistance (*cat*) | This study |

Supplementary Table 2 | Plasmids used in this study.

| **Plasmid** | **Characteristics** | **Reference** |
|---|---|---|
| pSK265 | Derivative form pC194; $cm^R$ | [4] |
| pSK265-*DsRed* | pSK265 expressing *DsRed* under the control of the *rpoB* promoter; $cm^R$ | [5] |
| pSK265-*sfGFP* | pSK265 expressing *sfGFP* under the control of the *rpoB* promoter; $cm^R$ | Lab Stock |
| pNP4 | pSK265 empty vector with *rpoB* promoter; $cm^R$ | This study |
| pNP7 | pSK265 expressing Timer under the control of the *rpoB* promoter; $cm^R$ | This study |
| pNP8 | pSK265 expressing Timer V105A under the control of the *rpoB* promoter; $cm^R$ | This study |
| pNP9 | pSK265 expressing $Timer^{FAST}$ under the control of the *rpoB* promoter; $cm^R$ | This study |

Supplementary Table 3 | Primers used in this study.

| **Primer** | **5' -> 3' Sequence** | **Reference** |
|---|---|---|
| NP61-DirMutS197T.FOR | TAG-TAC-CCT-GGT-AGC-TGC | This study |
| NP62-DirMutS197T.REV | CTA-TGT-TGA-CAC-CAA-ACT-GGA-TAT-AAC | This study |
| NP74-DirMutN42Q.FOR | CGA-AGG-CCA-CCA-GAC-CGT-AAA-GC | This study |
| NP75-DirMutN42Q.REV | TAT-GGC-CTC-CCC-TCT-CCT | This study |
| NP76-DirMutV105A.FOR | CGG-TGG-CGT-CGC-AAC-TGT-AAC | This study |
| NP77-DirMutV105A.REV | TCT-TCA-AAG-TTC-ATG-ACC-CTT-TCC-C | This study |

REFERENCES


[1] Peng HL, Novick RP, Kreiswirth B, Kornblum J, Schlievert P. Cloning, characterization, and sequencing of an accessory gene regulator (agr) in Staphylococcus aureus. J Bacteriol 1988;170:4365–72. https://doi.org/10.1128/jb.170.9.4365-4372.1988.

[2] Pohl K, Francois P, Stenz L, Schlink F, Geiger T, Herbert S, Goerke C, Schrenzel J, Wolz C. CodY in Staphylococcus aureus: a regulatory link between metabolism and virulence gene expression. J Bacteriol 2009;191:2953–63. https://doi.org/10.1128/JB.01492-08.

[3] Diep BA, Gill SR, Chang RF, Phan TH, Chen JH, Davidson MG, Lin F, Lin J, Carleton HA, Mongodin EF, Sensabaugh GF, Perdreau-Remington F. Complete genome sequence of USA300, an epidemic clone of community-acquired meticillin-resistant Staphylococcus aureus. Lancet 2006;367:731–9. https://doi.org/10.1016/S0140-6736(06)68231-7.

[4] Jones CL, Khan SA. Nucleotide sequence of the enterotoxin B gene from Staphylococcus aureus. J Bacteriol 1986;166:29–33.

[5] Caire R, Audoux E, Thomas M, Dalix E, Peyron A, Rodriguez K, Pordone N, Guillemot J, Dickerscheit Y, Marotte H, Vandenesch F, Laurent F, Josse J, Verhoeven PO. YAP promotes cell-autonomous immune responses to tackle intracellular Staphylococcus aureus in vitro. Nat Commun 2022;13:6995. https://doi.org/10.1038/s41467-022-34432-0.
